# Lamins organize the global three-dimensional genome from the nuclear periphery

**DOI:** 10.1101/211656

**Authors:** Xiaobin Zheng, Jiabiao Hu, Sibiao Yue, Lidya Kristiani, Miri Kim, Michael Sauria, James Taylor, Youngjo Kim, Yixian Zheng

## Abstract

Lamins are structural components of the nuclear lamina (NL) that regulate genome organization and gene expression, but the mechanism remains unclear. Using Hi-C, we show that lamins maintain proper interactions among the topologically associated chromatin domains (TADs) but not their overall architecture. Combining Hi-C with fluorescence *in situ* hybridization (FISH) and analyses of lamina-associated domains (LADs), we reveal that lamin loss causes expansion or detachment of specific LADs in mouse ES cells. The detached LADs disrupt 3D interactions of both LADs and interior chromatin. 4C and epigenome analyses further demonstrate that lamins maintain the active and repressive chromatin domains among different TADs. By combining these studies with transcriptome analyses, we found a significant correlation between transcription changes and the changes of active and inactive chromatin domain interactions. These findings provide a foundation to further study how the nuclear periphery impacts genome organization and transcription in development and NL-associated diseases.

**Highlights:**

1. Lamin loss does not affect the overall TAD structure but alters TAD-TAD interactions
2. Lamin null ES cells exhibit decondensation or detachment of specific LAD regions
3. Expansion and detachment of LADs can alter genome-wide 3D chromatin interactions
4. Altered chromatin domain interactions are correlated with altered transcription

## Introduction

The double nuclear membranes of the eukaryotic nucleus are perforated by thousands of nuclear pore complexes (NPCs) that mediate transport between the nucleus and the cytoplasm (Dickmanns et al., 2015). The outer and inner nuclear membranes are further bridged by the Linker of Nucleoskeleton and Cytoskeleton (LINC) complex, which is a physical link between nuclear envelope (NE) trans-membrane proteins (NETs) containing KASH and SUN domains and the cellular cytoskeleton (Wilson and Foisner, 2010). Beneath the inner nuclear membrane, the NPCs, SUN, additional NETs, and other proteins bind to a meshwork formed by the intermediate filament proteins known as lamins (Dechat et al., 2008; Gruenbaum and Foisner, 2015). This lamin-containing protein meshwork, often referred to as the nuclear lamina (NL), exists in all metazoan cell nuclei and is thought to play important housekeeping functions. Recent studies, however, have shown that NL proteins have developmental and tissue homeostatic functions (Barton et al., 2016; Chen et al., 2013; Chen et al., 2014; Kim et al., 2011; Razafsky et al., 2016; Young et al., 2012). Mutations in lamins or NETs also affect specific tissues in their associated human diseases. One partial explanation for this tissue specificity is the finding that lamins and NETs are often differentially expressed in different cell types (Czapiewski et al., 2016; Korfali et al., 2012). Despite the observed developmental defects of lamin-B1 and B2 double knockout (DKO) mice, mouse ES cells (mESCs) lacking these B-type lamins or all lamins (B1, -B2, and -A/C, triple knockout, TKO) are viable and can self-renew, similar to wild type (WT) (Kim et al., 2013b). Thus, at least some NL proteins are not required for housekeeping functions essential for cell viability but have roles in development.

Indeed, the B-type lamin, LAM, in *Drosophila*, functions in the cyst stem cells to support testis development by ensuring proper nuclear EGF signaling (Chen et al., 2013). Similarly, lamin-B1 in mammals regulates nuclear EGF signaling to promote dendrite development (Giacomini et al., 2016). The A-type lamins (lamin-A/C) and some NETs such as emerin can transduce mechano-signals into transcriptional responses (Ho et al., 2013; Qi et al., 2016). Although these and other studies have implicated NL proteins in connecting cell signaling to transcriptional regulation, it remains unknown how NL proteins could ensure that only specific genes among many potential target genes are activated by the signals.

The NL has been shown to create a repressive environment to silence its associated genes. Indeed, various NL proteins, including lamins, directly bind to transcription repressors that contribute to the formation of NL-associated heterochromatin (Wilson and Foisner, 2010). Tethering genes to the NL, however, does not always lead to their repression (Finlan et al., 2008; Kumaran and Spector, 2008; Reddy et al., 2008). Similarly, removing the major lamins, B1 and B2, expressed in mESCs does not lead to a global derepression of genes associated with the NL (Kim et al., 2011), but causes gene expression changes throughout the genome. Thus, how NL proteins could influence transcription by interacting with chromatin is poorly understood.

Applying the DNA adenine methyltransferase identification (DamID) technique to cultured mammalian cells has led to the mapping of NL-associated chromatin domains referred to as Lamina-Associated Domains, LADs (Guelen et al., 2008). By modeling chromatin landscapes based lamin-B1 DamID, histone modifications, core and linker histone enrichment in mESCs, we further separated LADs into two distinct chromatin domains defined as <u>Hi</u>stone and <u>L</u>amina l<u>ands</u>cape (HiLands)-B and -P, which together with four non-NL-associated HiLands-R, -O, -Y, and -G make up the six chromatin types of mESC genome (Zheng et al., 2015). The HiLands-B and -P divide LADs into distinct chromatin states with HiLands-P covering a longer stretch of chromatin and having higher lamin-B1 DamID values but lower H3K27me3 than HiLands-B (Zheng et al., 2015). The HiLands-B and -P defined by our HMM show good correlation, respectively, with the facultative and constitutive LADs inferred based on lamin-B1 DamID data from multiple cell types (Meuleman et al., 2013; Zheng et al., 2015).

Interestingly, lamin-B1-DamID studies in different cell lineages derived from ESCs reveal a cell-type specific detachment of facultative LADs (Peric-Hupkes et al., 2010) that correspond to HiLands-B (Zheng et al., 2015). Additionally, lamin-A/C and lamin-B receptor (LBR, a NET) are reported to sequentially tether heterochromatin to the NL, and their absence contributes to a large scale relocation of NL-associated heterochromatin to nuclear centers in cells such as the rod photoreceptors in nocturnal mammals (Solovei et al., 2013). These studies suggest that NL proteins play a role in three-dimensional (3D) genome re-organization important for development. How an NL protein regulates 3D chromatin interaction from the nuclear periphery remains, however, unknown.

If a given NL protein tethers some chromatin regions at the nuclear periphery, its absence may lead to the detachment of these chromatin from the NL, which could not only bring the genes in these chromatin regions into a different nuclear environment, but could also potentially disrupt interior chromatin interactions and global gene expression. Therefore, to understand how the NL influences 3D genome organization and transcription during development, it is essential to understand how the loss of NL proteins alters 3D chromatin interactions in relatively uniform progenitor cells that can differentiate into different cell types. Insights gained from this kind of analysis could help to further understand how 3D chromatin organization is regulated by the NL and how it impacts transcription throughout the genome during cell differentiation.

A high throughput chromosome conformation capture technique called Hi-C (Lieberman-Aiden et al., 2009) has enabled the study of 3D chromatin organization in human ESCs (hESCs) and mESCs (Dixon et al., 2012; Jin et al., 2013; Sexton et al., 2012). These studies suggest that the genome is organized into higher order structural domains referred to as <u>T</u>opologically <u>A</u>ssociated <u>D</u>omains (TADs). High resolution mapping of 3D chromatin organization within a TAD using 5C analyses reveal that interactions of the chromatin architectural proteins, such as CTCF, mediators, and cohesins, with chromatin result in formation of chromatin loops that further fold into sub-TADs and TADs (Dekker and Mirny, 2016; Nora et al., 2017; Phillips-Cremins et al., 2013; Sanborn et al., 2015).

Unlike CTCF and cohesins, however, NL proteins such as lamins exhibit neither sequence-specific DNA binding as seen in CTCF nor form rings/bracelets such as cohesins that produce chromatin loops. Instead, lamins form a dense meshwork concentrated at the nuclear periphery. The lamin meshwork may, therefore, regulate chromatin organization and transcription using different means from those employed by cohesins and CTCF. By combining Hi-C, 4C, and fluorescence *in situ* hybridization (FISH) with DamID, HiLands, epigenome, and transcriptome analyses in WT and lamin-null mESCs, we report that the lamin meshwork orchestrates 3D genome organization by differentially regulating different LADs, which in turn contributes to maintaining TAD-TAD interactions and interactions among chromatins with different activities.

## Results

### Lamin loss does not affect overall TAD structure, but alters inter-TAD interactions

To study the role of lamins in genome organization, we derived mESCs deleted of all three genes (*Lmnb1*, *Lmnb2* and *Lmna*) encoding all lamin isoforms to create lamin-TKO mESCs (Kim et al., 2011; Kim et al., 2013b). We performed Hi-C to map chromatin interactions in lamin-TKO and WT mESCs with the same genetic background and >87% euploidy (Kim et al., 2013b) using a ligation method similar to the *in situ* ligation method (Nagano et al., 2013; Rao et al., 2014). Our WT and TKO mESCs exhibited similar cell cycle profiles as judged by fluorescence activated cell sorting (FACS), indicating that the complete removal of lamins did not cause appreciable cell cycle alterations (Figure S1A). Two biological replicates of Hi-C data were generated for WT and TKO mESCs. After filtering and mapping, we obtained 304,631,864 and 374,862,869 total reads for WT and TKO mESCs, respectively (detailed statistics in Table S1). We processed the Hi-C data using Iterative Correction and Eigenvector decomposition (ICE) method (Imakaev et al., 2012) to obtain normalized contact maps for each Hi-C dataset. To assess the consistency of our Hi-C datasets, we analyzed chromatin-chromatin interactions (contact frequency) and plotted it against the linear genome distance between the pair of interacting sites. Similar contact frequency curves were found among the biological replicates (Figure S1B). Whole genome correlation efficient among WT and TKO replicates are also high at different genome window sizes (Figure S1C). Additionally, our data show low percentages of inter-chromosome reads, indicating low random ligation noise (Nagano et al., 2013; Rao et al., 2014) (Table S1). To further validate our Hi-C data, we compared our WT Hi-C datasets to the previously published WT E14 mESC Hi-C dataset (Dixon et al., 2012) and found a good consistency (Figure S1D).

To study whether lamin deletion affects TAD structure in mESCs, we used the Insulation Score method to call TADs from the Hi-C data. The Insulation Score is defined by the sum of interactions across each genomic window and local minima of Insulation Scores indicate TAD boundaries (Crane et al., 2015). We obtained 3,268 and 3,206 TAD boundaries from our WT and TKO mESCs, respectively (Table S2). Our WT TAD boundaries largely overlapped with those called from the WT E14 mESC Hi-C (Figure S1E).

The contact maps of an example region show highly similar TAD structures in our WT and lamin-TKO mESCs (Figure 1A-B). Direct comparison of the TAD boundaries also reveals >90% overlap between our WT and TKO mESCs (Figure 1C). By plotting the Insulation Score profiles around the TAD boundary, we found highly similar patterns for both the overlapping and non-overlapping TAD boundaries (Figure 1D). This suggests that the small number of non-overlapping TAD boundaries are due to minor random variations of Insulation Scores. Thus, Hi-C studies show that deletion of all lamins does not disrupt the overall TAD structure in mESCs.

**Figure 1.**
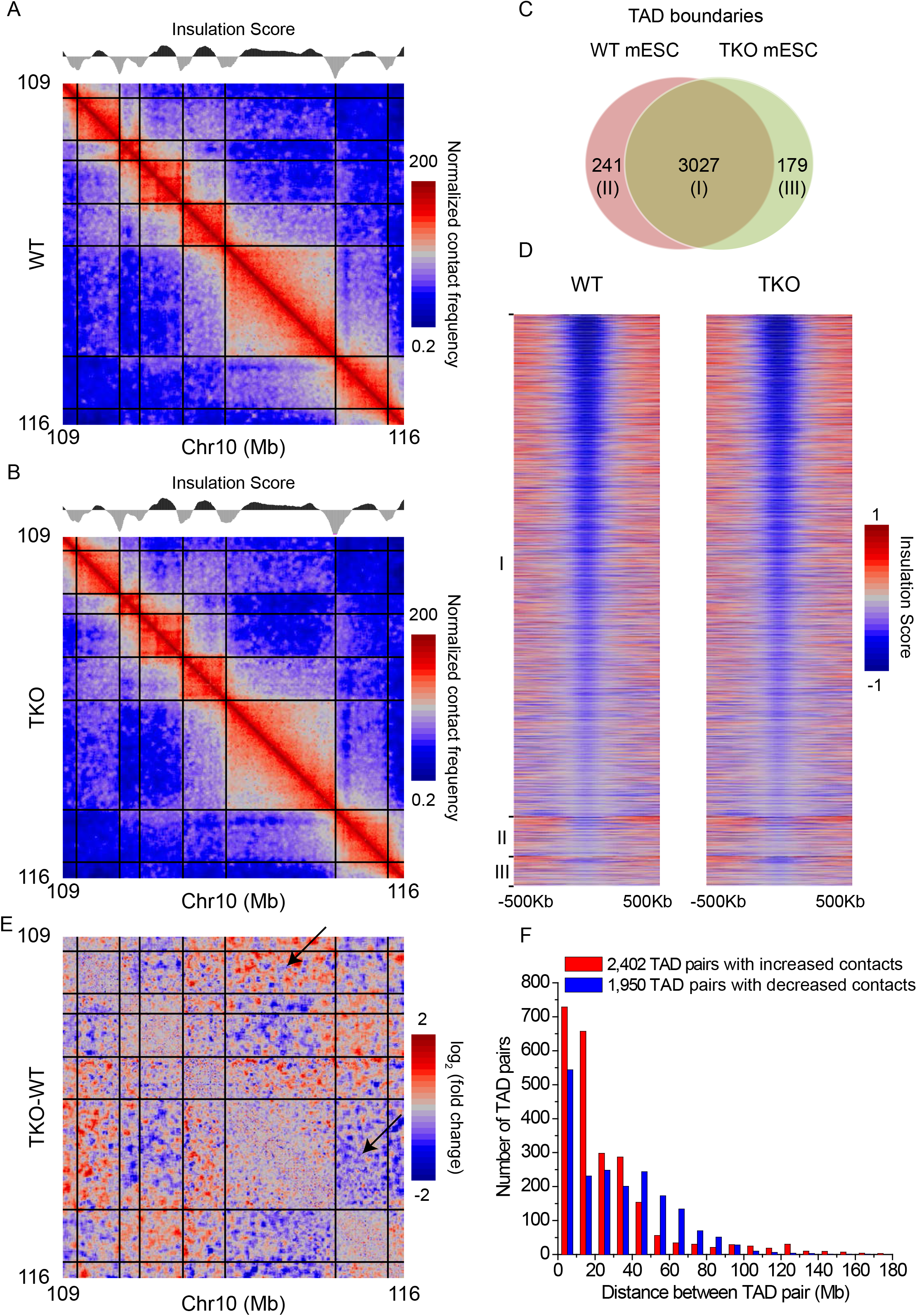
Effects of lamin deletion on TADs in mESCs. **A-B**. Dynamic-binned heatmaps showing the normalized contact frequencies and TAD structures in a representative region on chr10 in WT (A) and lamin TKO (B) mESCs at 20-Kb resolution. Each bin is expanded to contain a minimum of 20 reads. Normalized contact frequency is calculated using the Iterative Correction and Eigenvector decomposition (ICE) method. The total sequencing depth in TKO is normalized to the same sequencing depth as the WT. The plots of Insulation Score along the same region are shown above the heatmaps. Local minimum values of Insulation Score indicate TAD boundaries, which are delineated by black lines in the heatmaps. **C.** Venn diagram showing the TAD boundary overlap between WT and lamin TKO mESCs. **D.** Heatmaps showing the insulation scores in ±500 Kb around TAD boundaries at 20-Kb resolution. I, II, and III indicate the three groups of TAD boundaries shown in C in WT and TKO mESCs. **E.** Dynamic-binned heatmaps showing the log_2_ fold change of interactions between TADs in the same representative chromosome 10 region shown in A and B at 20 Kb resolution. Each bin is expanded to contain a minimum of 20 reads. Black lines delineate TAD boundaries. The black arrows indicate examples of TAD pairs showing increased or decreased interactions. **F.** Histogram showing the distributions of inter-TAD distances of TAD pairs exhibiting altered interactions upon lamin loss.

Further comparison of Hi-C interactions between our WT and lamin-TKO mESCs, however, did reveal clear changes of TAD-TAD interactions upon lamin loss (Figure 1E). Using edgeR to identify TAD pairs exhibiting altered interactions in our WT-TKO comparisons, we found 4,352 TAD pairs exhibiting altered interactions (False Discovery Rate, FDR<0.05) upon lamin loss (Table S2). To further validate these TAD-TAD interaction changes, we analyzed whether the altered TAD-TAD interactions found in our WT-TKO comparisons are also altered in the E14 WT-TKO comparisons. We found that 96.2% of the increased or 91.8% of the decreased TAD-TAD interactions observed in our WT-TKO comparisons were recapitulated in the E14 WT-TKO comparisons (Figure S1F). Thus TAD-TAD interaction changes are not due to random variation or genetic background differences but are caused by lamin deletion. Plotting the TAD-TAD interaction changes against the distances between the TAD pairs, we found that the increased and decreased interactions have different distance distributions (Figure 1F). Thus, lamin loss does not affect overall TADs organizations, but alters TAD-TAD interactions in mESCs.

### Lamin loss differentially affects the interaction of chromatin domains in TADs

To understand how lamin loss affects the interactions of chromatin domains, we first analyzed whether and how the TAD-TAD interaction changes correlated with LADs defined by lamin-B1 DamID (Peric-Hupkes et al., 2010). For all the TAD pairs exhibiting altered interactions upon lamin loss in mESCs, we analyzed the average lamin-B1 DamID values in each TAD. We found that both TADs in a TAD pair exhibiting increased interactions had high positive DamID values (Figure 2A), whereas the TAD pairs with decreased interactions often involved one TAD with high positive DamID values and the other TAD with negative DamID values (Figure 2B). This indicates that lamin deletion differentially affects inter-TAD interactions within the NL and between the NL and nuclear interior chromatin.

**Figure 2.**
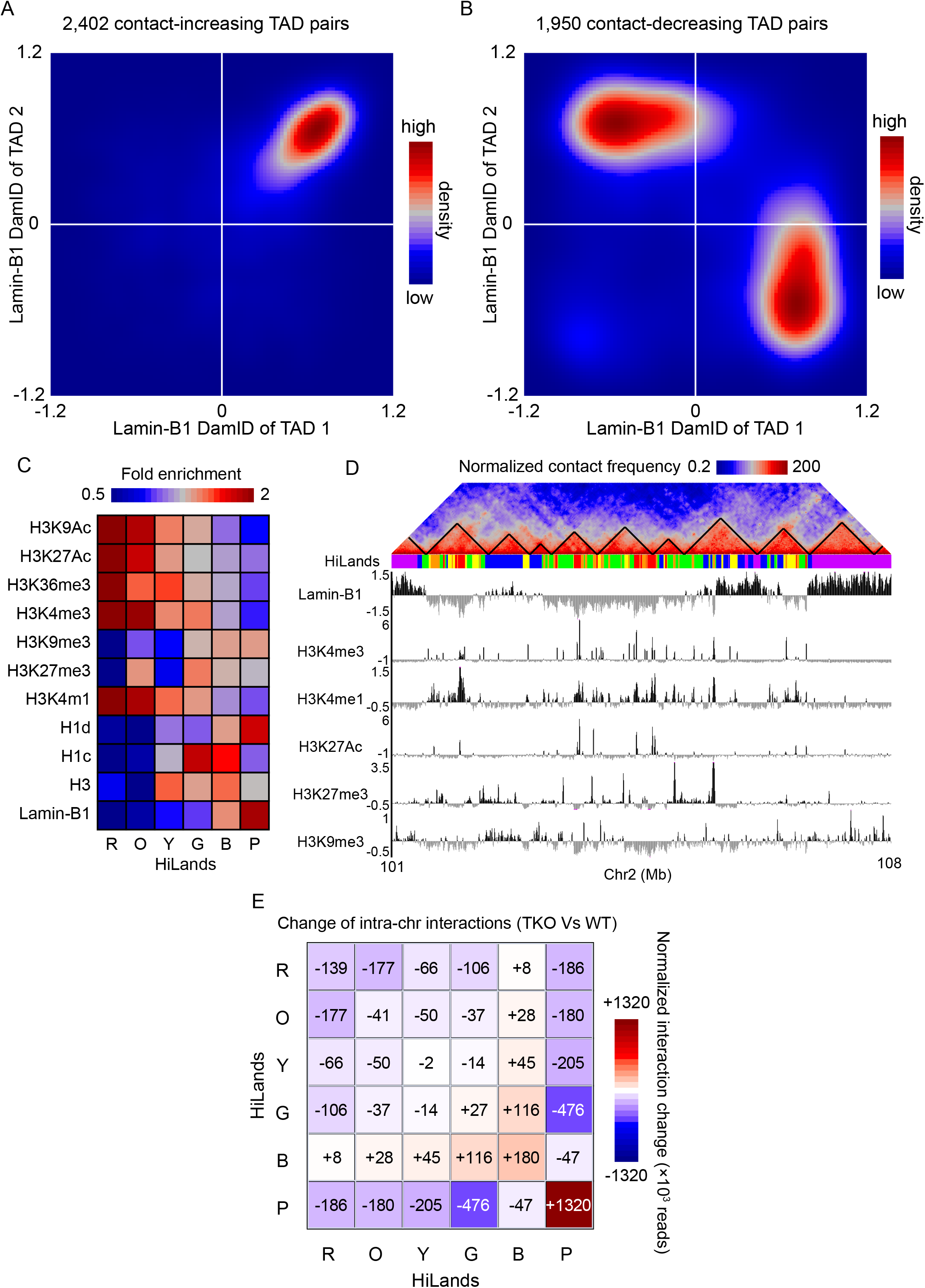
Lamins regulate global 3D genome interactions among chromatin domains with different features. **A-B.** Contour plots showing the number of TAD pairs having increased (A) or decreased (B) interactions upon lamin loss as a function of the lamin-B1 DamID values of the two TADs. **C.** Heatmap showing the enrichment of different histone modifications, histone H1 and H3, and lamin-B1 DamID reads on the six different HiLands in mESCs. H3, H1c, and H1d shown are fold enrichment normalized against input. Histone modifications shown are fold enrichment against H3. **D.** An example region on chromosome 2 showing Hi-C heatmap with TADs delineated by black lines, color coded HiLands, lamin-B1 DamID, and histone modifications. The Hi-C heatmap is dynamic-binned and normalized as in Figure 1. Histone modifications shown are log_2_ fold enrichment against H3. **E.** Heatmaps showing the normalized interaction change of total intra-chromosome interactions within each HiLands and between HiLands pairs upon lamin loss in mESC. The numbers of total intra-chromosome interactions in TKO were first normalized to the same as those in WT. Then the normalized numbers of total intra-chromosome interactions within and between HiLands in TKO were subtracted by the corresponding numbers of interactions in WT to get the changes. The numbers of increased (+) or decreased (-) interaction changes are shown.

To understand how lamin loss influences both LADs and interior chromatin, we utilized a chromatin state model that we developed previously using a Hidden Markov Model (HMM) of 7 markers: LADs (lamin-B1 DamID), the enrichment of histones (H3, H1c, and H1d), and histone modifications H3K4me1, H3K27me3, and H3K9me3 (Zheng et al., 2015). We established six chromatin states in mESCs referred to as <u>Hi</u>stone-<u>L</u>amina l<u>ands</u>capes (HiLands)-Red (R), Orange (O), Yellow (Y), Green (G), Blue (B), and Purple (P). As shown in Figure 2C and D, four of these HiLands (HiLands-R, O, Y, and G) delineate interior chromatin regions (low lamin-B1 DamID values) with HiLands-R exhibiting the highest transcriptional activity (Zheng et al., 2015). The remaining two HiLands (-B and -P) divide LADs into distinct chromatin states with HiLands-P covering a longer stretch of chromatin and having higher lamin-B1 DamID values but lower H3K27me3 than HiLands-B (Figure 2C and D) (Zheng et al., 2015). HiLands-B and -P correspond to the facultative and constitutive LADs defined using different cell types (Zheng et al., 2015). Depending on the genome location, a TAD can contain multiple different HiLands, while some long HiLands-P can contain more than one TAD (Figure 2D). When mapping the Hi-C interactions to HiLands in WT mESCs, we found distinct interactions both within and between different HiLands. WT mESCs exhibit strong intra-chromosome interactions within all HiLands and between HiLands pairs R-O, Y-G, and B-P, whereas the interactions between HiLands-B or P and the interior HiLands are low (Figure S2A).

When comparing the HiLands interactions between WT and lamin-TKO mESCs, the strongest increase in intra-chromosome interactions occur within HiLands-P followed by -B and -G and between HiLands-B and interior HiLands, whereas the interactions between HiLands-P and the other HiLands are decreased in the TKO mESCs (Figure 2E). There are also decreased intra-chromosome interactions within and between most of the interior HiLands upon lamin loss (Figure. 2E). By analyzing the inter-TAD interaction changes in the context of HiLands, we found that the increased interactions within HiLands-P and decreased interactions between HiLands-P and the other HiLands (see Figure 2E) account for the majority of intra-chromosomal TAD-TAD interaction changes (Figure S2B-C) as observed in Figure 2A-B.Thus, lamins play a context-specific role in maintaining peripheral chromatin architecture, which ensures proper global chromatin interactions in mESCs.

### Lamin loss causes de-condensation of LADs characterized by HiLands-P

To understand whether lamin loss causes physical changes of chromatin, we next studied LADs characterized by HiLands-P which have higher lamin-B1 DamID values than HiLands-B (see Figure 2C and D). We first asked at what linear chromatin length scale HiLands-P chromatin exhibits strongest interactions with itself by plotting the log_2_ fold change of interactions upon lamin loss against the distance between the two interacting loci in HiLands-P. We found a moderate but clear decrease in interactions between two loci in HiLands-P at <1 Mb length scale, but the interactions increased from ~1 Mb with the strongest increase occurring at 10-20 Mb upon lamin loss (Figure 3A). Since 1 Mb is a typical length of TAD in HiLands-P (Figure S3A), most of the increase in interactions in HiLands-P may occur between different TADs. Indeed, by counting the total number of altered interactions within HiLands-P chromatin, we found an increased inter- and decreased intra-TAD interactions upon lamin loss (Figure S3B).

**Figure 3.**
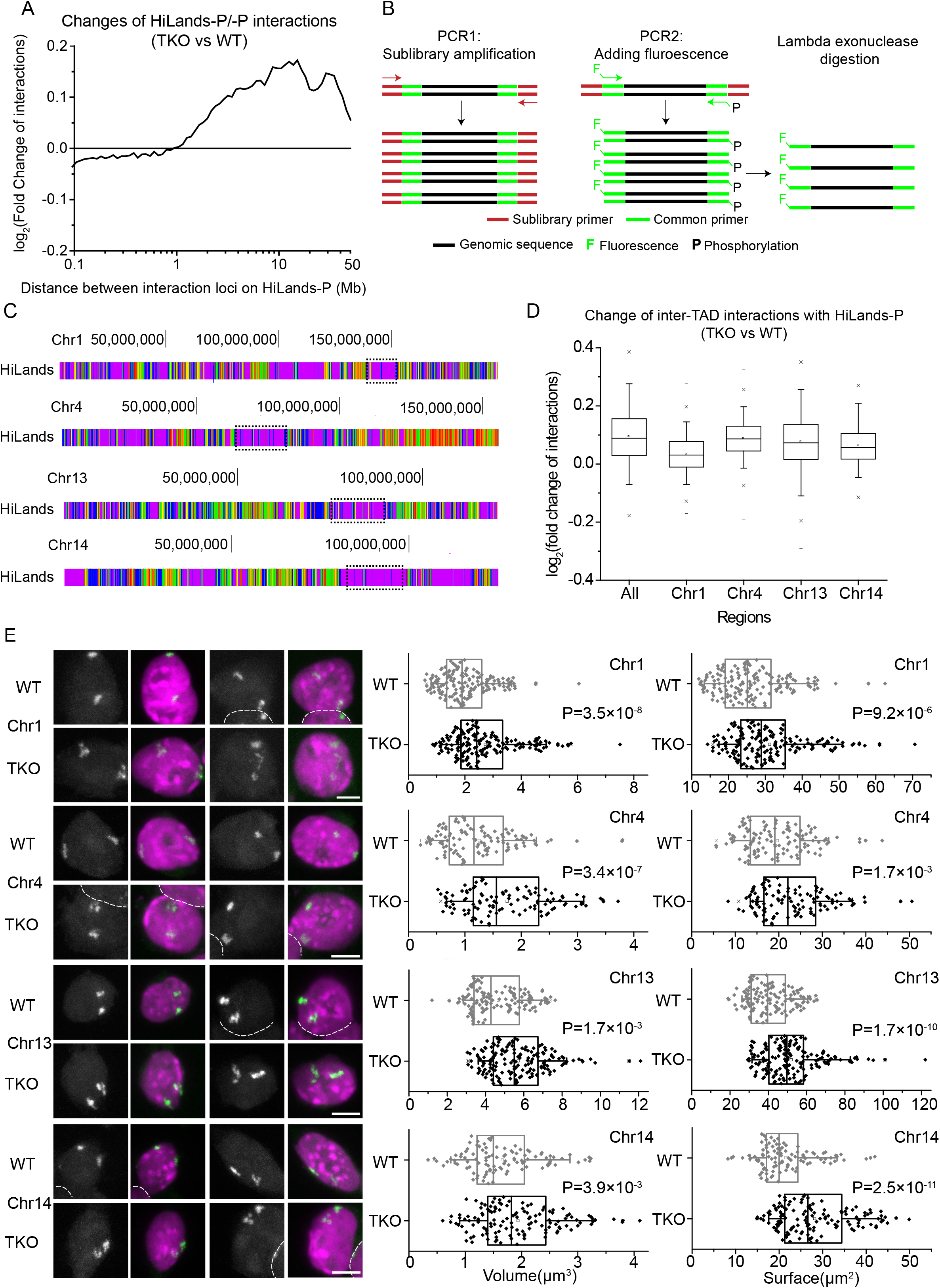
Expansion of HiLands-P upon lamin loss. **A.** A plot of log_2_ fold increased or decreased total interactions between two HiLands-P regions upon lamin loss as a function of the distance between the regions. **B.** FISH probe production. PCR1 amplifies the probes for a specific sub-library using the indicated sub-library primers. PCR2 produces the labeled sub-library probes using the fluorescently labeled and phosphorylated common primers. Lambda exonuclease digests the phosphorylated DNA strand to produce the single stranded DNA probes for FISH. **C.** Four regions (dashed boxes) on Chromosome 1, 4, 13, and 14 consisting of mostly HiLands-P were selected for FISH. HiLands are shown in corresponding colors. **D.** Box plot showing the log_2_ fold change of inter-TAD interactions for 20-Kb windows in the whole genome (All) or in selected chromosome regions shown in C. Only HiLands-P interactions are included. **E.** Two representative 3D-projection FISH images for each of the four selected regions in C. Purple: DAPI staining for DNA. White: FISH signal. The white dashed lines demarcate the boundaries of nuclei that are next to one another. Scale bars, 5 µm. The volume and surface areas of the four chromatin regions are quantified to the right. P-values, Wilcoxon rank-sum test.

The increase in inter-TAD HiLands-P interactions could be caused by a physical decondensation of HiLands-P chromatin upon lamin loss. To visualize whether HiLands-P chromatin is decondensed, we used fluorescence *in situ* hybridization (FISH). Based on the Oligopaint method (Beliveau et al., 2012), we designed densely arrayed oligos for four HiLands-P-enriched regions on chromosomes 1, 4, 13, and 14. Probes for FISH were generated by two rounds of PCR (Figure 3B). The first set of PCR primers used in round 1 had specific sequences that allowed the amplification of a pool of oligos (sub-library) corresponding to the selected regions (Figure 3C). The second round of PCR used a common primer pair shared by all sub-libraries that allowed fluorescence labeling of each amplified sub-library (Figure 3B, see Table S3 for the primer sequences). The central 50-bp genomic sequences were selected based on a published algorithm (Rouillard et al., 2003). The regions selected had different levels of inter-TAD interaction increases in HiLands-P (Figure 3D). Using double-blind measurements of the FISH signal based on the ‘Spots object’ function in Imaris, we found that both the volumes and surface areas of these regions increased significantly in TKO mESCs compared to WT (Figure 3E, S3C). Since a previous study in a human colon cancer cell line, DLD-1, showed that lamin-B1 depletion by RNAi resulted in decondensation of the entire chromosomes 18 and 19 by FISH (Camps et al., 2014), we performed Oligopaint using oligo probes made from the entire chromosome 1 or 13 Oligopaint libraries and found no significant differences in either the volumes or surface areas of the two chromosomes between TKO and WT mESCs (Figure S3D). This difference could be due to cell type differences or lamins may play a stronger role in maintaining chromatin condensation in differentiated cells than in the ESCs. Thus, lamin loss in mESCs results in decondensation of HiLands-P LADs without appreciably affecting the compactness of whole chromosomes.

### Lamin loss causes compartment shift of LADs characterized by the detachment of HiLands-B from the NL

We next analyzed the impact of lamin loss on HiLands-B LADs that exhibit lower lamin-B1 DamID values than HiLands-P LADs (see Figure 2C and D). Previously, we have used emerin DamID to study how lamin-B1 and -B2 gene knockout (lamin-B DKO) in mESCs affected the interaction between LADs and the NL and found that HiLands-B and P LADs exhibited a decreased and increased emerin DamID values, respectively, upon the loss of the two B-type lamins (Zheng et al., 2015). However, another emerin DamID study, using the same DKO mESCs or the DKO mESCs depleted of lamin-A/C (equivalent to lamin TKO mESCs), argued that lamin loss did not affect the overall interaction between LADs and nuclear periphery (Amendola and van Steensel, 2015). Considering that mESCs express a very low level of lamin-A/C (Eckersley-Maslin et al., 2013), it is expected that the lamin-B1/B2 DKO and lamin TKO mESCs should have similar emerin DamID maps. Indeed, our analyses of the published emerin-DamID datasets of the lamin TKO and WT mESCs (Amendola and van Steensel, 2015) revealed a decrease in emerin-DamID score in >75% HiLands-B LADs regions and a score increase in >75% HiLands-P LADs regions upon lamin loss (Figure 4A, Table S4), similar to those of the lamin-B DKO mESCs we reported (Zheng et al., 2015). We believe that our separation of LADs into HiLands-B and P has increased the power of analyses, which has led to the identification of the differential regulation of LADs by lamins that was previously missed by Amendola and Steensel.

**Figure 4.**
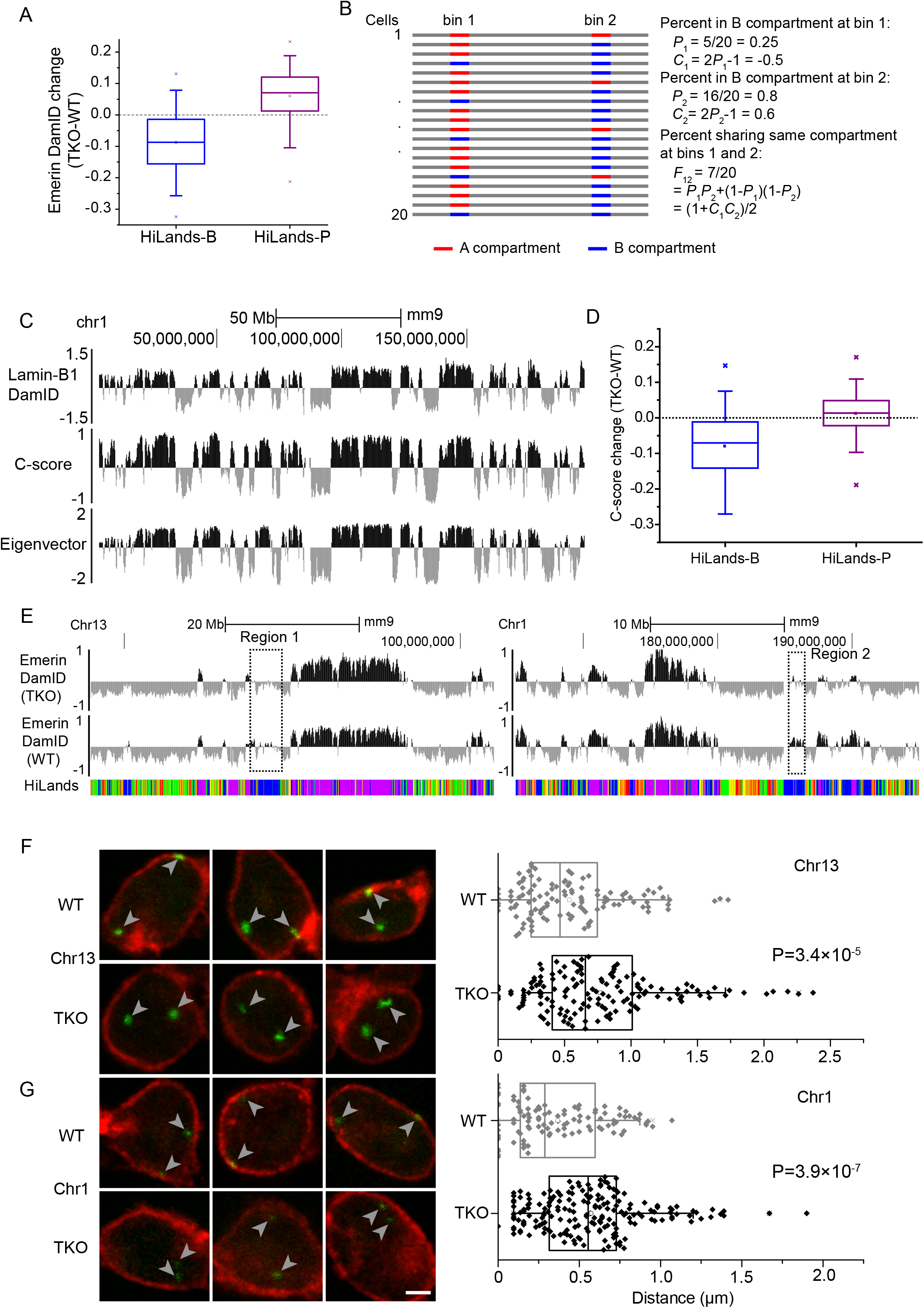
Lamin loss causes HiLands-B detachment from the NL. **A.** Box plot showing the changes of emerin-DamID values on HiLands-B and -P throughout the genome upon lamin loss in mESCs. **B.** Definition of C-score. *P*_1_ or *P*_2_ represents the percentage of cells in which genomic windows 1 or 2 (bin 1 or 2), respectively, is in the B compartment. *C*_1_ and *C*_2_ represent the C-scores of bins 1 and 2 and they have a linear relationship to *P*_1_ and *P*_2_, respectively. *F*_12_ is the percentage of cells with both bin 1 and 2 in the same compartment. **C.** Genome-browser views showing a good consistency of lamin-B1 DamID values, C-scores, and eigenvector values along chromosome 1 in WT mESCs. The eigenvector is calculated using the Homer software at 20-Kb resolution. For the C-score and eigenvector, positive and negative values indicate B and A compartment, respectively. **D.** Box plot showing the changes of C-scores in HiLands-B and -P throughout the genome upon lamin loss in mESCs. **E.** Genome browser views of emerin-DamID in WT and lamin-TKO mESCs on Chr 1 (right) and 13 (left). The dashed boxes highlight the regions 1 and 2 used for producing FISH probes, which correspond to the HiLands-B enriched chromatin with reduced emerin-DamID values upon lamin loss in mESCs. Grey arrowheads point to the FISH signals. **F-G.** Three sets of representative images of emerin immunostaining (red) and FISH (green) of region 1 (Chr13, F) and region 2 (Chr1, G) are shown to the left. Each image is a plane from a confocal stack. The shorter length of the region 2 compared to 1 resulted in weaker FISH signals seen in **G**. Scale bars, 2 μm. Quantification of the distances from each FISH signal to the nuclear membrane (emerin) are shown to the right. P-values, Wilcoxon rank-sum test.

Next, we used our Hi-C data to further verify the reduced interaction of HiLands-B LADs with the nuclear periphery detected based on DamID. Previous Hi-C studies have separated the 3D genome into A and B compartments because higher interactions are found within each compartment than between the two (Lieberman-Aiden et al., 2009). Further analyses revealed that the A compartment is gene-rich, high in H3K36me3 and DNaseI sensitive regions, whereas the B compartment is gene-poor with relatively low transcriptional activity, corresponding to LADs consisting of HiLands-B and P (Lieberman-Aiden et al., 2009). Since HiLands-B exhibit reduced emerin DamID values upon lamin loss (Figure 4A), we reasoned that these chromatin regions may shift their localization away from the B compartment whereas HiLands-P should mostly remain in the B compartment. If this shift does occur, it should be detectable by analyzing the compartment changes of chromatin using our Hi-C datasets. Unfortunately, the original compartment analysis was based on the first interaction matrix eigenvector, which detected Hi-C interaction differences between the A and B compartments within one experimental sample (Lieberman-Aiden et al., 2009), and is thus unsuitable for quantitative comparisons between different Hi-C experiments. To allow quantitative comparisons, it is important to develop a model to describe the probability that in an individual cell a genomic region (bin) is in either the A or B compartment. We defined the compartment score (C-score) as a linear function of the probability (P) so that the percentage (F) of cells in which two bins were in the same compartment had a linear relationship to the product of the C-scores (C) in the two bins (Figure 4B). We then used a maximum-likelihood method to infer the C-scores for each genomic bin. Briefly, for two windows *i* and *j*, we have the probability (*F*_*ij*_) that windows *i* and *j* are in the same compartment*F*_*ij*_ = (1 + *C*_*i*_*C*_*j*_)/2. Since in individual cells, two windows in different compartments are unlikely to interact with each other, the number of Hi-C contacts between two genomic windows *i* and *j* should be proportional to *F*_*ij*_. By including the effect of linear genomic distance and experimental biases, we built a Poisson model *n*_*ij*_ ~ *Poisson*(*B*_*i*_*B*_*j*_*H*(*d*_*ij*_)(1 + *C*_*i*_*C*_*j*_)) where *n*_*ij*_ is the number of observed contacts between windows *i* and *j*, *d*_*ij*_ is the linear distance, *H*(*d*_*ij*_) is the function of the distance dependency, *B*_*i*_ and *B*_*j*_ are the bin bias factors. This model leads to a log-likelihood function ln 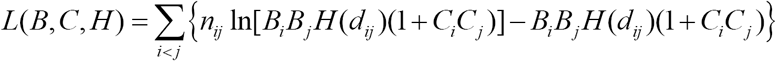, which allows us to estimate all the parameters including the C-scores by iterative optimization (see Supplementary methods for details).

Our calculated C-scores have positive and negative values corresponding to B and A compartments, respectively. We found a good consistency of C-scores between the biological Hi-C replicates of our WT mESCs and between our WT and published WT E14 mESCs Hi-C datasets (Figure S4A). The positive and negative C-scores are also highly correlated with the positive and negative lamin-B1 DamID values, respectively, in WT mESC (Figure S4B). The A and B compartments inferred in our WT mESCs by C-scores and by the eigenvector method are also highly consistent with the B compartment having high positive lamin-B1 DamID values (Figure 4C). Thus, the C-score can be used to quantitatively compare genomic loci in A or B compartments across different experiments.

When applying the C-score analysis to our WT and TKO mESCs, we found that over 75% of HiLands-B regions had decreased C-scores, upon lamin loss, which was highly consistent with the result from our emerin DamID data (Figure 4A and D). Moreover, the C-score and emerin-DamID changes on HiLands-B show a good genome wide correlation (Figure S4C). We only observed a small C-score increase of HiLands-P upon lamin loss which is consistent with these chromatin regions remain in compartment B. The increased interaction between HiLands-P and nuclear periphery as measured by emerin DamID is likely caused by the decondensation of HiLands-P LADs. Therefore, two different techniques, DamID and Hi-C, demonstrate that lamin loss results in HiLands-B LADs chromatin exhibiting loss or reduced interactions with the nuclear periphery and moving from compartment B to A.

To visualize if HiLands-B LADs indeed move away from the NL upon lamin loss, we selected two regions consisting of mostly HiLands-B and exhibiting decreased emerin DamID values upon lamin loss (Figure 4E) for FISH analyses using Oligopaint. The levels of emerin-DamID reduction in the selected regions 1 and 2 with respect to the genome-wide HiLands-B reduction upon lamin loss are shown in Figure S4D. To quantify the distance between the FISH signal and the NL, the ‘Cells object’ function in Imaris was used to create the surface of all nuclei defined by emerin staining, whereas the ‘Spots object’ function marked the FISH signals. The distance from the center of each FISH signal and the nearest nuclear surface was measured in the ‘Cells object’. We found that the two HiLands-B regions in TKO mESCs were farther away from the NL than in the WT (Figure 4F-G, S4E).

We also measured the volumes and surface areas of the two HiLands-B regions and found that they were similar in both TKO and WT mESCs (Figure S4F). The HiLands-B LADs were smaller than HiLands-P LADs, which could account for our inability to detect HiLands-B decondensation by FISH in TKO mESCs. Alternatively, H3K27me3 is enriched on HiLands-B LADs, but depleted on HiLands-P LADs (Zheng et al., 2015) (see Figure 2C-D). Since the polycomb complex 1 (PRC1) can bind to H3K27me3 to cause chromatin condensation (Boettiger et al., 2016; Simon and Kingston, 2009), lamin loss may not affect HiLands-B LADs condensation. Nonetheless, lamins ensure NL localization of LADs characterized by HiLands-B.

### Lamins influence transcription by regulating 3D chromatin domain interactions

The altered 3D chromatin interactions upon lamin loss shown above may affect the epigenome in the lamin TKO mESCs. We first analyzed the heterochromatin markers H3K9me3 and H3K27me3 in TKO and WT mESCs by ChIP-seq. There were no significant overall changes of these histone modifications between WT and TKO mESCs (Table S5). This is different from the observations in fibroblasts harboring lamin-A mutation, which exhibit a significant overall change in these heterochromatin modifications and in 3D chromatin interactions upon senescence (McCord et al., 2013).

Next, we analyzed the transcriptome differences between lamin-TKO and WT mESCs by ribosome-RNA-depleted RNA-seq. The TKO mESCs had 385 up- and 841 down-regulated genes compared to WT (Figure 5A, Table S6). Consistent with our studies of lamin-B DKO mESCs (Kim et al., 2011), we found no enrichment of differential expression of NL-associated genes (Figure 5B). Instead, the differentially expressed genes are distributed across all six HiLands (Figure S5A). Moreover, we found only 3 genes within the detached HiLands-B were up regulated (Table S6). This is consistent with previous findings that nuclear periphery localization of genes does not necessarily lead to their repression (Finlan et al., 2008; Kumaran and Spector, 2008; Reddy et al., 2008)

**Figure 5.**
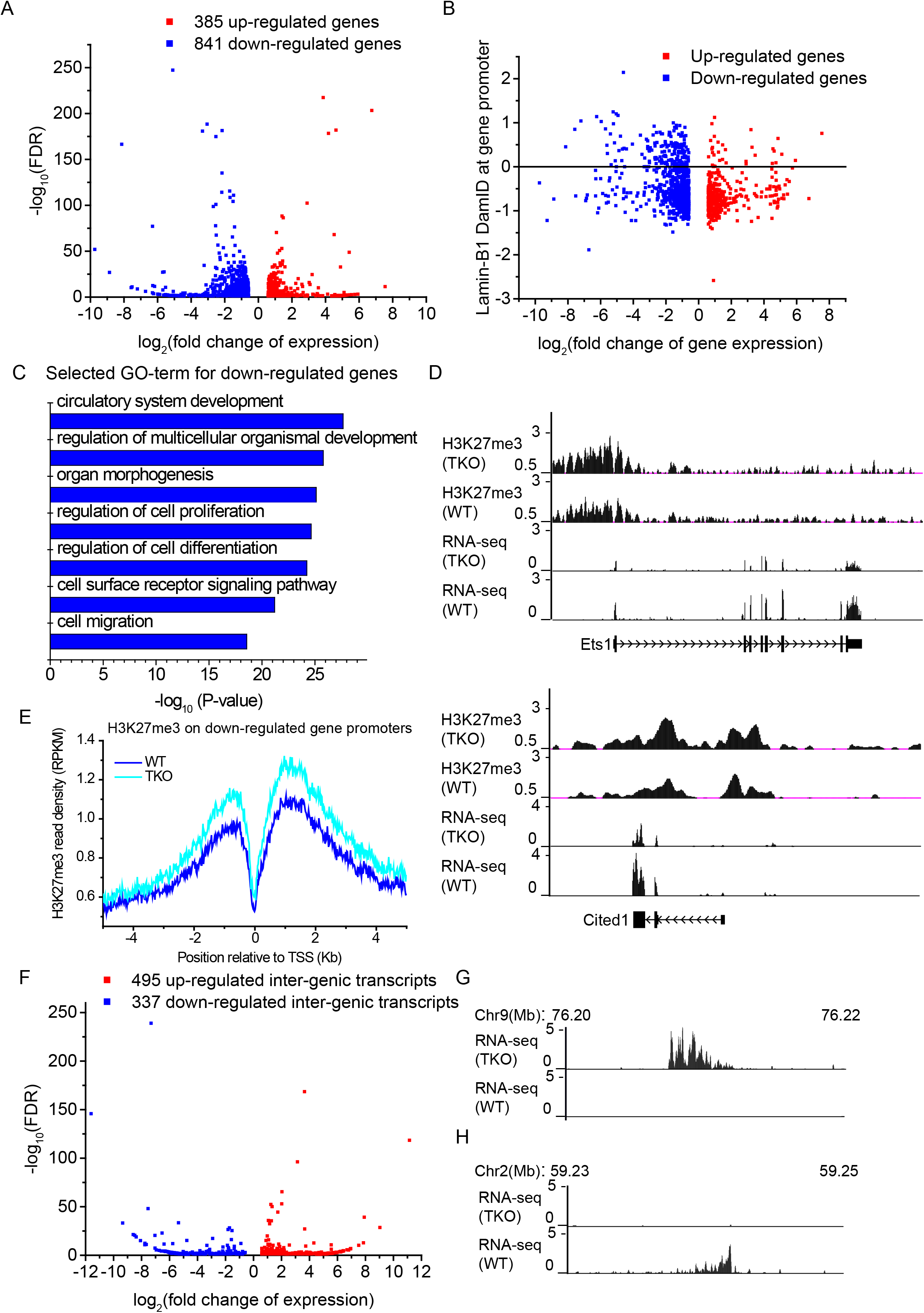
The effect of lamin loss on transcription in mESCs. **A.** Volcano plot of RNA-seq data showing the gene expression change and the statistical significance. Threshold of differential expression: fold change>1.5, FDR<0.05. **B.** Scatter plot showing a lack of enrichment of altered genes in LADs (based on lamin-B1 DamID in WT mESCs. P=0.9, hypergeometric test) upon lamin loss in mESCs. **C.** The top significant GO terms of the down-regulated genes upon lamin loss in mESCs. **D.** The genome browser views showing the H3K27me3 and RNA-seq tracks on two down-regulated genes upon lamin loss in mESCs that are required for development. **E.** Distribution of H3K27me3 reads within 5 Kb up- and down-stream of the down-regulated gene promoters. H3K27me3 showed a significant increase upon lamin loss on these promoters (P<2×10^−16^, Wilcoxon signed-rank test). **F.** Volcano plot of RNA-seq data showing the expression change of unannotated inter-genic transcripts and the statistical significance. Threshold of differential expression, fold change>1.5, FDR<0.05. **G-H.** The genome browser views showing the up- (**G**) or down-regulated (**H**) non-genic transcripts upon lamin loss. Genomic positions of the loci in mm9 are shown at the top.

Gene Ontology (GO) analyses revealed an enrichment of down-regulated genes with functions in differentiation, development, and morphogenesis (Figure 5C, Table S6) but no significant GO term for up-regulated genes. Although there were no genome-wide changes for H3K27me3 and H3K9me3 levels, when analyzing the promoter regions of the down regulated genes we found that H3K27me3, but not H3K9me3, exhibited significant up regulation upon lamin loss (Figure 5D-E, S5B). Interestingly, we recently found that lamin TKO mice were lethal between embryonic day (E)9.5 and E13.5 (data not shown). The increased H3K27me3 on the down-regulated genes upon lamin loss could disrupt their timely expression thereby contributing to the developmental failure. We also observed altered expression of intergenic regions distributed across all six HiLands upon lamin loss in mESCs (Figure 5F-H, S5C, Table S6). ChIP-seq of H3K4me3 showed that the alteration of this active histone mark on the transcriptional start sites (TSS) correlated with the transcriptional changes of both annotated genes and intergenic transcripts upon lamin loss in mESCs (Figure S5D).

Since lamin TKO mESCs exhibited HiLands-P decondensation and HiLands-B detachment, which leads to the altered 3D chromatin interactions among TADs containing different HiLands, we analyzed whether these changes are correlated with gene expression. By plotting the number of changed genic and intergenic transcripts in HiLands-R, O, Y, and G against the distance of the transcription start sites (TSS) to their nearest HiLands-B on the linear genome, we found that the closer the TSS were to HiLands-B, the more likely their transcription would change in TKO mESCs (Figure 6A). Importantly, the TSS of both up- and down-regulated transcripts are distributed significantly closer to HiLands-B than those of all transcripts (Figure S6A). Thus, the change in 3D chromatin interactions due to HiLands-B detachment is correlated with an increased chance of expression changes of the nearby transcripts.

**Figure 6.**
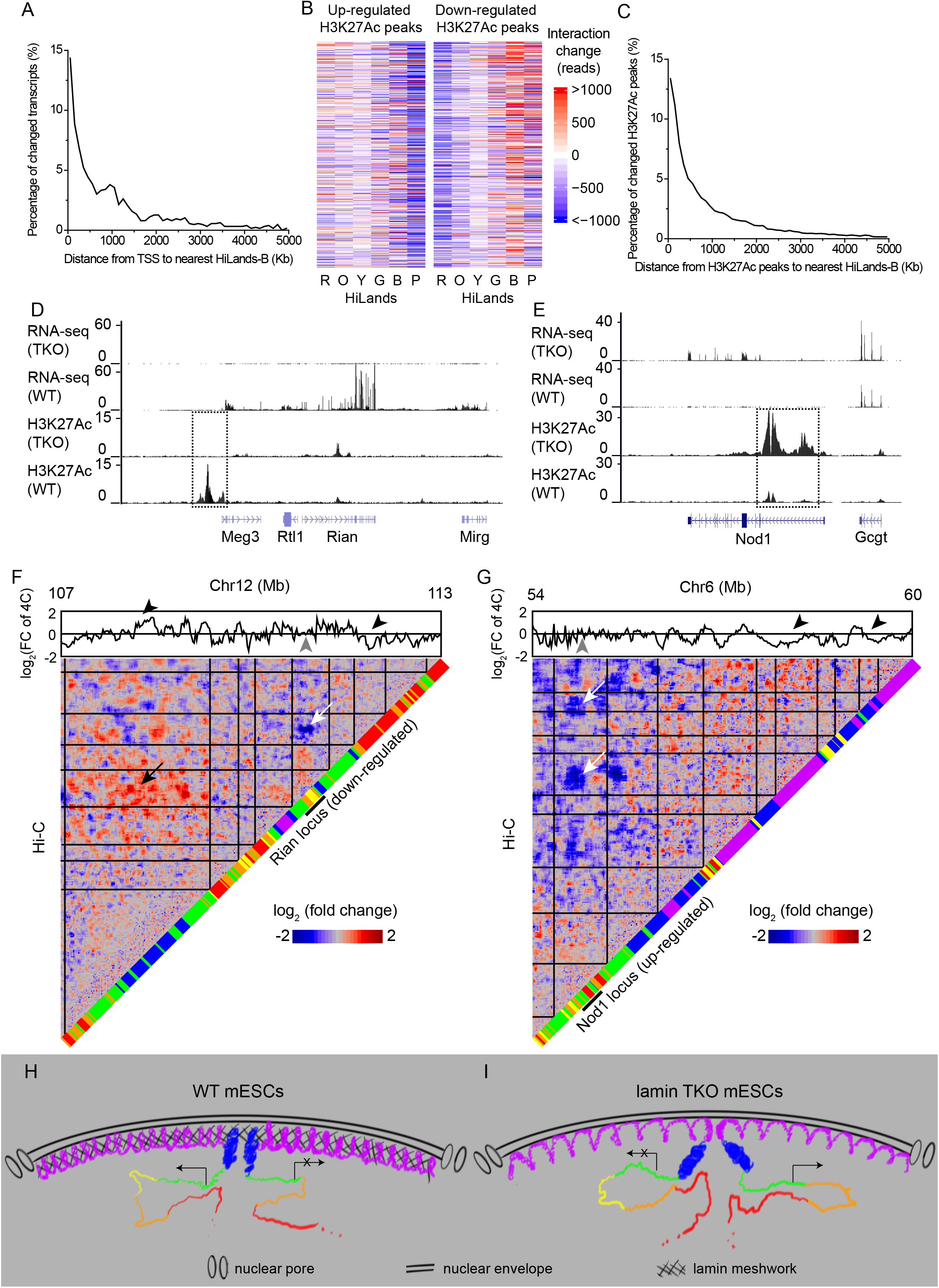
Effects of 3D chromatin interaction changes on transcription upon lamin loss. **A.** A plot showing the distance distribution of transcription start sites (TSS) of altered transcripts upon lamin loss to the border of the nearest HiLands-B. Only the TSS in HiLands-R, -O, -Y and -G are plotted. **B.** Heatmaps showing that H3K27Ac increased peaks have decreased interactions with HiLands-P (P<2×10^−16^, Wilcoxon signed-rank test), whereas H3K27Ac decreased peaks have decreased interaction with HiLands-R (P<2×10^−16^, Wilcoxon signed-rank test) and increased interaction with HiLands-B (P<2×10^−16^, Wilcoxon signed-rank test) in the 100Kb genomic windows surrounding each peak upon lamin loss. **C.** A plot showing the distance distribution of the up- or down-regulated H3K27Ac peaks to the nearest HiLands-B upon lamin loss in mESCs. **D-E.** The down- (**D**) and up-regulated (**E**) expressions at Rian and Nod1 loci upon lamin loss (top two tracks), which correlate with the changes of enhancer activities (marked by dashed boxes) as indicated by the H3K27Ac (bottom two tracks). **F-G.** 3D genome interaction changes around the Rian (**F**) and Nod1 (**G**) loci. The heatmaps show the change of Hi-C interactions using the same setting as in Figure 1E with the black lines marking TADs. HiLands are shown along the diagonal with the loci indicated by the short black lines. Change of 4C interactions are shown above the heatmaps (grey arrowheads indicate the 4C baits). The white and black arrows in F indicate the decreased and increased interactions of the Rian locus with Hilands-R and HiLands-B, respectively, in different TADs. The white arrows in G indicate the decreased interactions of the Nod1 locus with HiLands-B and -P in different TADs. The black arrowheads indicate the corresponding regions showing increased or decreased interactions with the bait region. The Hi-C and 4C interaction changes show significant linear correlation: Rian, P<2×10^−16^; Nod1, P=5×10^−13^, t-test. **H-I.** A lamin meshwork-cage model. Loss of the lamin meshwork in the WT mESC nucleus (**H**) leads to the expansion of HiLands-P, which dislodges HiLands-B from the NL in lamin TKO mESCs (**I**). The detachment of HiLands-B in turn disrupts the active and inactive chromatin neighborhoods, thereby leading to global transcriptional changes. The six HiLands are color coded. The expression and repression of genes are indicated.

Changes in 3D chromatin interactions in TKO mESCs could alter the interactions between active and inactive chromatin domains, leading to changes in enhancer activities and altered transcription. Indeed, ChIP-seq of H3K27Ac (an active enhancer marker) and H3K4me1 (a marker for all enhancers) in TKO and WT mESCs showed that H3K4me1 did not show significant changes (Table S5), whereas H3K27Ac exhibited changes on many chromatin regions (Table S5). We found that the up-regulated H3K27Ac peaks had reduced interactions with the transcriptionally repressive HiLands-P, whereas the down-regulated H3K27Ac peaks exhibited increased interactions with the transcriptionally repressive HiLands-B and decreased interactions with the transcriptionally active HiLands-R (Figure 6B). Plotting the number of changed H3K27Ac peaks against their distance to the nearest HiLands-B showed that the closer the enhancers were to HiLands-B the more likely they would change their activities (Figure 6C). Importantly, the up- and down-regulated H3K27Ac peaks are distributed significantly closer to HiLands-B than those of all H3K27Ac peaks (Figure S6B). Additionally, the increased and decreased H3K27Ac are also correlated with the increased and decreased transcription, respectively, upon lamin loss in mESCs (Figure S6C). We also analyzed the distances from the TSS of the altered transcripts and the altered H3K27Ac peaks to Hilands-P and found that they are further away from HiLands-P than from HiLands-B (Figure S6D and E). Thus, the altered transcription and enhancer activities upon lamin loss are more related to HiLands-B detachment than to HiLands-P expansion.

To test whether gaining the interaction of non-LADs regions with detached HiLands-B at the TADs scale are related to transcription changes, we analyzed the 1,609 TADs in non-LADs regions and found 176 showed significantly increased (but not decreased) interactions with at least one TAD containing ≥80% HiLands-B. Of these 176 TADs, 70 (39.8%) have at least one down-regulated gene. By contrast, of the remaining 1,433 TADs in non-LADs regions, only 190 (13.3%) showed at least one down-regulated gene. Thus the nuclear interior TADs that gained interactions with TADs containing ≥80% HiLands-B upon lamin loss exhibited an enrichment of down-regulated genes (39.8% compared to13.3%, P=2×10^−8^, hypergeometric test).

To further analyze the relationship between 3D chromatin interaction and transcription changes, we selected a down- (Rian) and an up-regulated (Nod1) gene cluster whose expression changes correlated with their changes of H3K27Ac (Figure 6D-E). We performed 4C analyses of the two loci in WT and TKO mESCs and plotted the 4C interaction changes against the Hi-C interaction changes and the corresponding HiLands types (Figure 6F-G). Consistent with the above global analyses, the down-regulation of the Rian gene cluster is correlated with its increased interactions with a TAD enriched for inactive HiLands-B and decreased interactions with a TAD enriched for active HiLands-R (Figure 6F). On the other hand, the up-regulation of the Nod1 gene cluster is correlated with its decreased interactions with two TADs enriched for inactive HiLands-P and B (Figure 6G). Thus, lamins can regulate transcription by controlling the condensation of HiLands-P and localization of HiLands-B at the NL to prevent inappropriate global interactions between chromatin domains with different transcriptional activities (Figure 6H-I).

## Discussion

### Differential impact of the lamin meshwork on the NL-associated chromatin

Although the mechanism by which the NL regulates chromatin has long been studied and interpreted in the conceptual framework of chromatin tethering by NL proteins (Gonzalez-Sandoval et al., 2015), how this tether influences global 3D chromatin organization remains unknown. We show that lamins do not regulate TADs, but are required for maintaining proper inter- and intra-TAD interactions. Our FISH studies reveal the detachment of HiLands-B from the NL and decondensation of HiLands-P at the NL upon lamin loss. These changes of different LAD regions are unexpected because HiLands-P have stronger interactions with the NL than HiLands-B based on DamID studies (Peric-Hupkes et al., 2010) and the tether model would predict a stronger detachment of HiLands-P from the NL than HiLands-B. Therefore, our findings demonstrate that the lamin meshwork does not regulate LADs as a simple tether for chromatin.

Our Hi-C analyses revealed an increased inter-TAD and decreased intra-TAD interactions among HiLands-P along the linear chromatin length upon lamin loss. The decondensed TADs within HiLands-P likely contribute to the increased inter-TAD interactions and HiLands-P expansion we detected by FISH. In contrast to HiLands-P, we did not detect clear HiLands-B decondensation. The smaller HiLands-B domain sizes relative to HiLands-P (Zheng et al., 2015) resulted in our use of FISH probes covering shorter genomic regions (<5 Mb) for FISH compared to HiLands-P (>12 Mb), which may not allow the detection of a small chromatin decondensation in HiLands-B. Alternatively, HiLands-B may not be decondensed upon lamin loss because these LADs regions are enriched for H3K27me3 (Zheng et al., 2015) and the polycomb proteins could help maintain chromatin condensation in lamin TKO mESCs.

How does lamin loss cause HiLands-P decondensation and HiLands-B detachment? Considering that lamins assemble into a dense filamentous meshwork at the NL, we propose a meshwork caging model that can explain all of our observations made by different genome techniques. Whereas other NL proteins may tether LADs chromatin, the lamin meshes function to trap and cage HiLands-P and, to a lesser degree, HiLands-B, to ensure their condensation, proper organization and position at the NL (Figure 6H). Since HiLands-B have weaker interactions with the NL than HiLands-P (DamID studies) and since HiLands-P comprise >60% of all LADs (Zheng et al., 2015), the decondensation of HiLands-P upon lamin loss could ‘push’ HiLands-B away from the NL (Figure 6I). The detached HiLands-B would in turn have an increased opportunity to interact with the less condensed interior chromatin (Figure 6I), thereby causing the increase in the intra- and inter-chromosome interactions between HiLands-B and other interior chromatin domains as detected by Hi-C in TKO mESCs. Although purified lamins exhibit weak interactions with histones and DNA (Wilson and Foisner, 2010), the dense lamin meshwork could support extensive multivalent interactions with LADs. This, coupled with the lamin-associated proteins that recognize DNA, histones, and modified histones, would allow the lamin meshwork to cage LADs to prevent the decondensation of HiLands-P and the detachment of HiLands-B.

### The influence of the lamin meshwork on 3D genome organization and transcription

Our studies shed light on how the 3D chromatin organization changes upon lamin loss could affect transcription of genic and intergenic regions globally in mESCs. We found that decondensation of HiLands-P and detachment of HiLands-B correlated with a disruption of existing chromatin domain interactions and created new ones in TKO mESCs. These chromatin interaction changes were further correlated with transcription changes. We did not find global up-regulation of lamin-bound genes upon lamin loss, which is consistent with previous finding that lamins do not generally repress genes at the NL (Amendola and van Steensel, 2015; Kim et al., 2011; Kim et al., 2013b). Additionally, detachment of HiLands-B from the NL does not lead to overall activation of genes in these LADs, either. This could be due to the enrichment of H3K27me3 on HiLands-B, which could silence genes independent of association with the NL. Importantly, we found that the gain of active enhancers correlated with increased interactions with active chromatin regions and gene expression, whereas loss of active enhancers correlated with increased interactions with inactive chromatin regions and lowered gene expression upon lamin loss. Thus, altered 3D chromatin interactions upon lamin loss can disrupt active or inactive chromatin neighborhoods, which in turn could impact transcription in both the NL-associated and interior chromatin (Figure 6H-I). Our findings provide a rationale for how lamins can function at the NL to globally influence transcription, a question that has perplexed the field for decades.

Our studies have focused on mESCs, but we believe that the concept revealed here should apply to differentiated cell types. The altered expression of many genes required for development in TKO mESCs reported here suggests that lamins could be required for their proper regulation during differentiation. By maintaining existing 3D chromatin interactions and helping to establish new ones, lamins may facilitate expression of additional genes critical for development. For example, lamins may establish and/or maintain 3D chromatin configurations to allow the activation of a subset of target genes in response to mechano-transduction or other chemical signals during development. Our TKO and WT mESCs should allow further dissection of how lamins can regulate differentiation as 3D chromatin organizers.

### A possible role for lamins in coupling morphogenesis to transcriptional regulation

Interestingly, the facultative LADs (fLADs), corresponding to HiLands-B, undergo differential detachment from the NL in different cell lineages (Meuleman et al., 2013; Zheng et al., 2015). Our findings suggest that such lineage-specific detachment would establish new 3D chromatin interactions, which could contribute to gene regulation during differentiation. Cell differentiation is accompanied by increased cytoskeleton activities and cell morphological changes (Lecuit and Lenne, 2007; Vong et al., 2010). Since the cytoskeleton is connected to the nucleus by LINC complexes and NPCs via lamins, it is tempting to speculate that a local cytoskeletal change might reduce the meshwork density of lamins near the altered cytoskeleton, thereby leading to decondensation and detachment of selected HiLands-P and HiLands-B, respectively. As the timing and type of cytoskeletal changes are specific to differentiating cell types, studying cytoskeleton- and lamin-regulated changes of LADs could potentially help to understand how morphological changes during development might impact global 3D genome organization and transcriptional wiring important for morphogenesis and organogenesis.

## STAR*METHODS

Detailed methods are provided in the online version of this paper and include the following:

-Key resources table
-Contact for reagent and resource sharing
-Experimental model mESCs and culture conditions
-Method details

1. 3D FISH, mESC FACS, and karyotyping
2. Hi-C library preparation and sequencing
3. ChIP-sequencing
4. Hi-C data analyses
5. 4C data analyses
6. DamID data analyses
7. RNA-seq data analyses
8. ChIP-seq data analyses

## Author contributions

Conceptualization, X.Z. and Y.Z.; Validation, X.Z., J.H., and Y.K.; Formal analyses, X.Z., J.H., Y.K., M.S., J.T., and Y.Z.; Investigation, X.Z., J.H., Y.K., S.Y., L.K., M.K., and Y.Z.; Writing—Original Draft, X.Z. and Y.Z.; Writing—review & editing, X.Z., J.H., Y.K., S.Y., L.K., M.K., M.S., J.T., and Y.Z.; Visualization, J.H. and X.Z. Bioinformatics, X.Z., M.S., J.T.; Supervision, Y.Z. and Y.K.

## Acknowledgements

We thank Allison Pinder and Fred Tan for help with deep sequencing, Mahmud Siddiqi for Imaris analyses, Ona Martin and Lynne Hugendubler for technical assistance, Robert Johnston and Kayla Viets for help with FISH, Mohammad Heydarian for suggestions on RNA-seq analyses, Joseph Tran, Chen-Ming Fan, Ona Martin, and Lynne Hugendubler for proofreading, and the Zheng lab for helpful discussions. Supported by the National Research Foundation of Korea (grant number: NRF-2014R1A1A1037106) and the Soonchunhyang University Research Fund to Y.K., and by GM056312, GM106023, and Ellison Medical Foundation to Y.Z.

## Supplementary Figures and Tables

**Figure S1.**
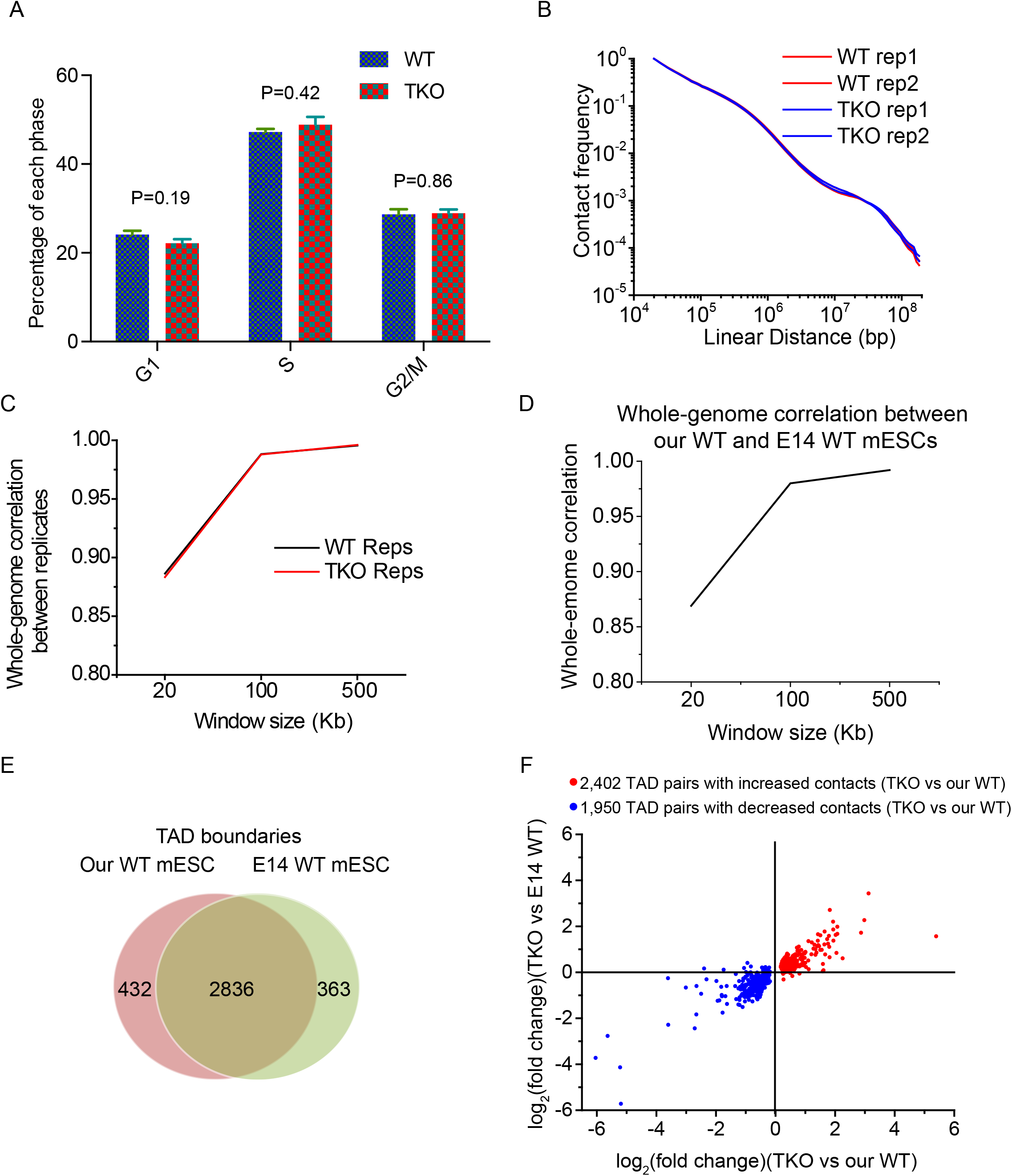
Validation and reproducibility analyses of our Hi-C studies (support Figure 1) **A.** Distribution of cell cycle phases. The percentages of cells in G1, S, and G2/M phases of the cell cycle were measured for our WT and lamin TKO mESCs by FACS. P-values, t-test. **B.** Contact frequencies between genomic regions as a function of linear genomic distance in our WT and lamin TKO replicates. All distance curves were normalized against the values at 20 Kb. **C.** Whole-genome correlation of normalized Hi-C interactions between WT or TKO replicates at 20-, 100- and 500-Kb resolutions using the Pearson correlation coefficient. **D.** Whole-genome correlation of normalized Hi-C interactions between our WT mESC Hi-C data (combining two replicates) and previously published WT E14 mESC Hi-C data (combining the published three replicates) (Dixon et al., 2012) at 20-, 100- and 500-Kb resolutions using the Pearson correlation coefficient. **E.** Venn Diagram showing the overlap of TAD boundaries called from our WT mESC Hi-C data and previously published E14 mESC Hi-C data. **F.** Comparison of inter-TAD interaction changes between our lamin TKO mESCs and our WT or previously published WT E14 mESC Hi-C data.

**Figure S2.**
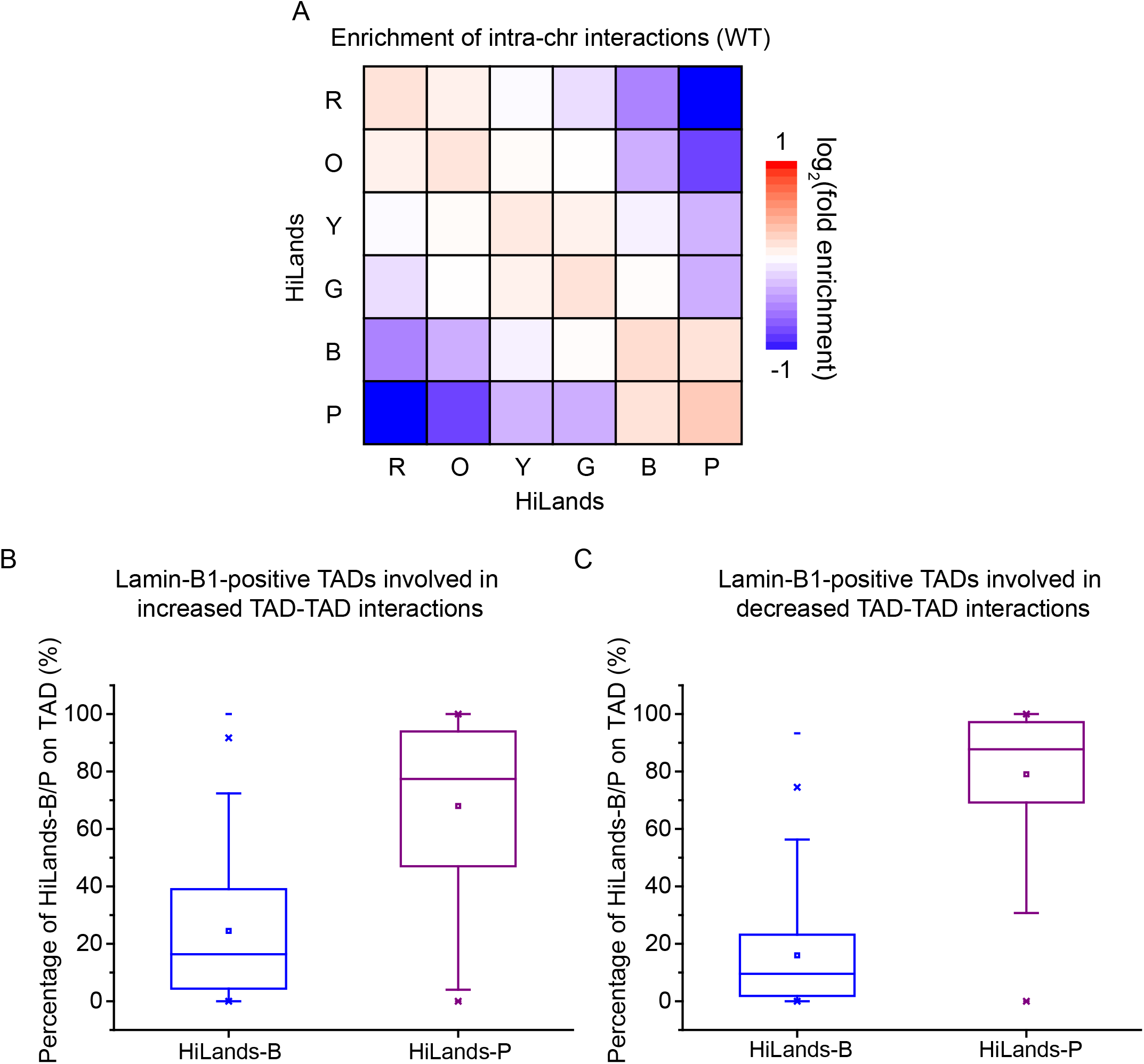
Mapping of Hi-C interactions to HiLands (Support figure 2) **A.** Heatmap showing the enrichment of intra-chromosome interactions in our WT mESC Hi-C data within and between the six HiLands. **B-C**. Box-plots showing the percentages of HiLands-B and P within the lamin-B1-positive TADs involved in increased (**B**, corresponding to Figure 2A) or decreased (**C**, corresponding to Figure 2B) TAD-TAD interactions upon lamin loss. In both cases, the TADs are covered more by HiLands-P than by HiLands-B (P<2×10^−16^, Wilcoxon signed rank test).

**Figure S3.**
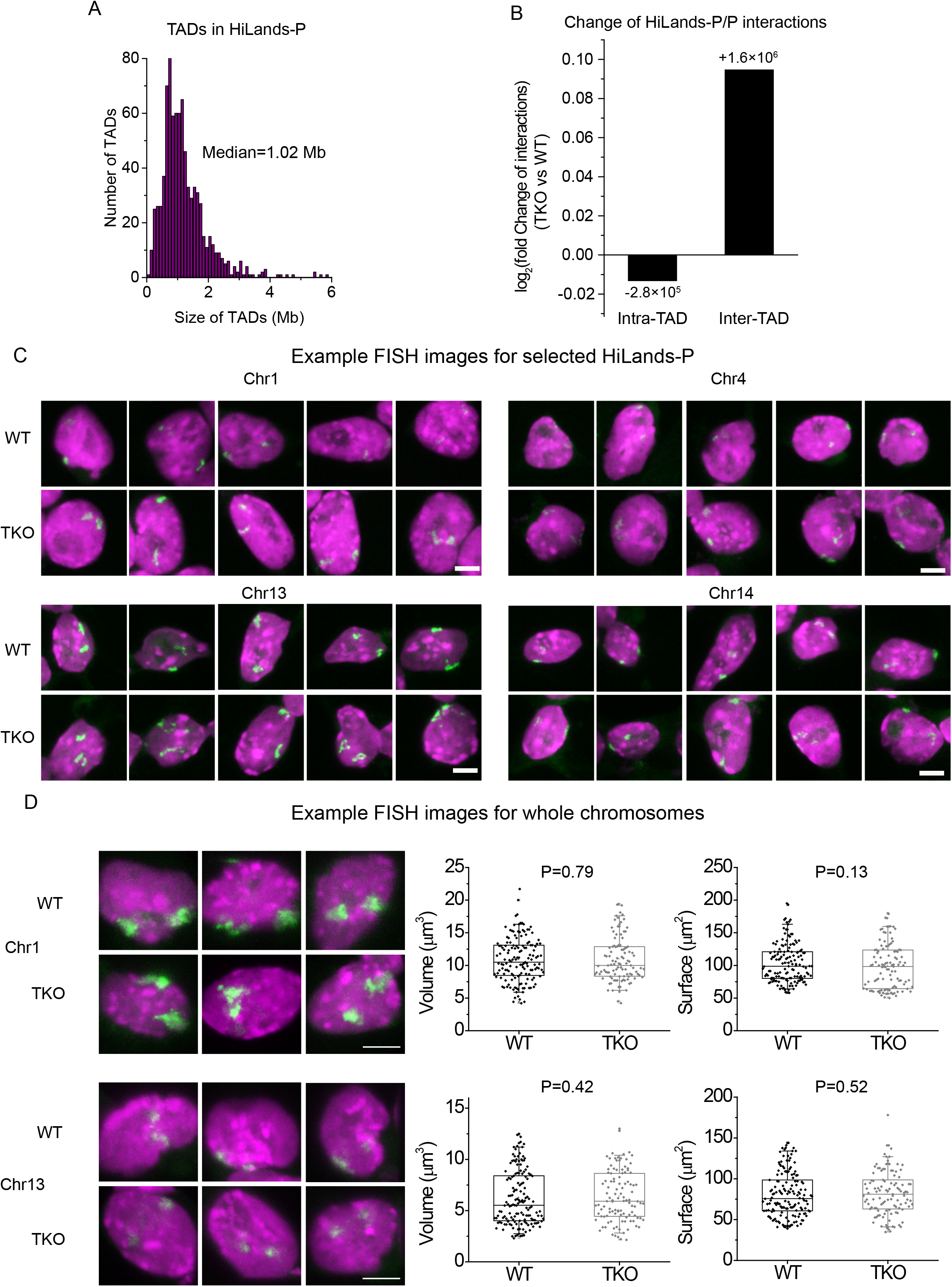
Analyses of HiLands-P TADs and FISH (support Figure 3) **A.** The distribution of the size of TADs in HiLands-P. **B.** Bar-plot showing the total log_2_ fold increased inter- or decreased intra-TAD interactions in HiLands-P. The number of normalized interaction change of total intra- or inter-chromosome interactions are shown. The normalization was the same as Figure 2E. **C.** Additional representative FISH images of HiLands-P on the indicated chromosomes in WT and TKO mESCs. **D.** Representative 3D projection images of FISH (green) of whole chromosomes 1 (Chr1) and 13 (Chr13) in our WT and lamin TKO mESCs. The nuclei were counter stained by DAPI (purple). The quantifications of the volume and surface area of these chromosomes are shown to the right of the images as dot plots. Scale bar, 5 μm.

**Figure S4.**
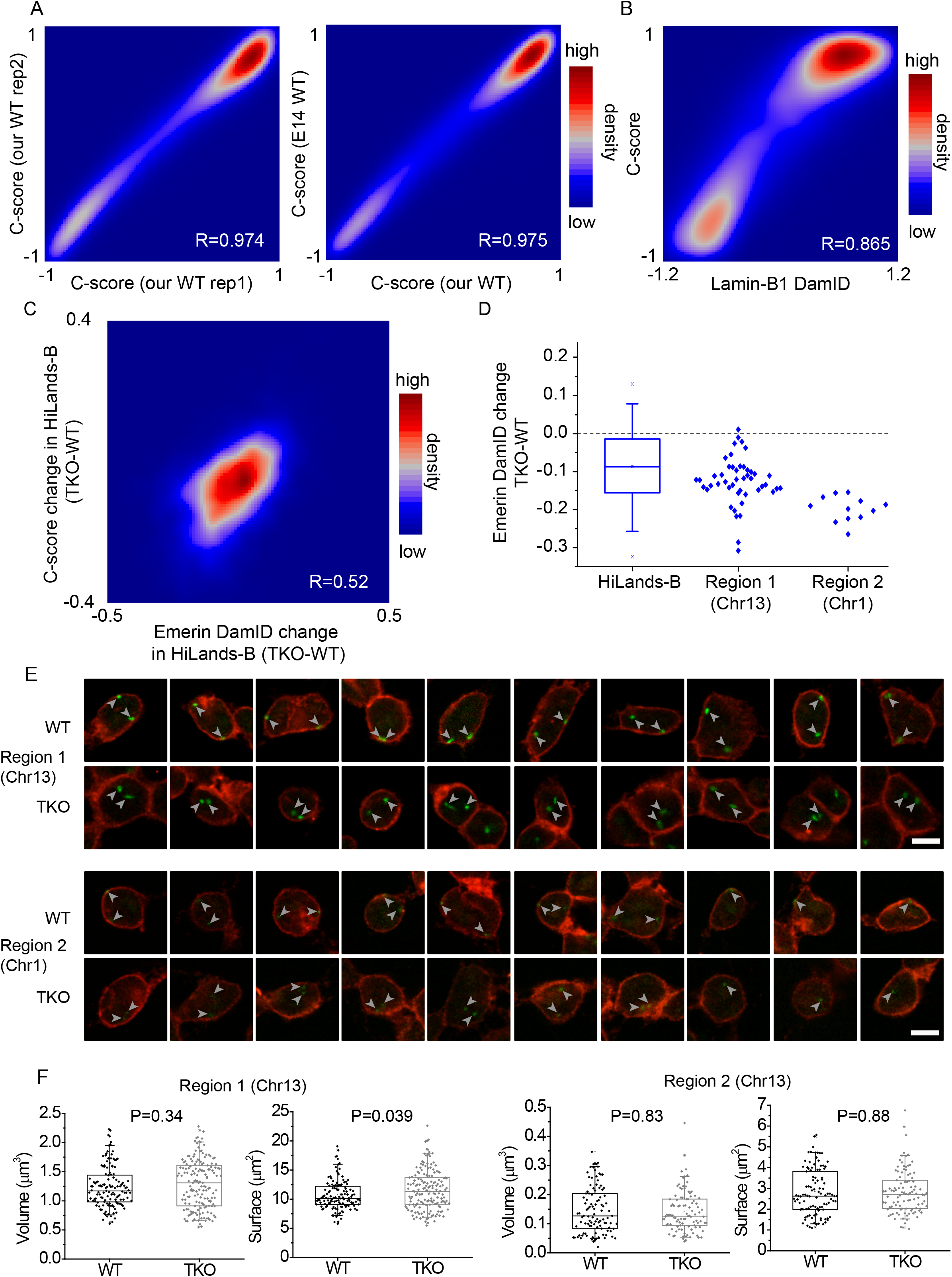
C-scores and FISH analyses (support Figure 4) **A.** Contour plots showing the correlation of C-scores between our two WT mESC replicates and between our WT and the published WT E14 mESC data at 20-Kb resolution. **B.** Contour plots showing the correlation between C-scores and lamin-B1 DamID values on the whole genome at 20-Kb resolution in WT mESCs. **C.** Contour plots showing the correlation between C-score change and emerin-DamID value change upon lamin loss on HiLands-B at 100-Kb resolution. **D.** The box plot showing the range of genome-wide reduction of emerin-DamID values in HiLands-B upon lamin loss in mESCs (left). The dot plots on the right show the range of reduction of emerin-DamID values within the highlighted regions 1 and 2 in Figure 4E. **E.** Additional FISH images of HiLands-B on the indicated chromosomes from WT and TKO mESCs. The last three set of images of Chr1 region 2 only show one FISH signal in the plane selected. The other FISH signals are outside of this plane. Grey arrowheads point to the green FISH signals. **F.** Dot plot quantifications of the volume and surface area of the 3D FISH of the two HiLands-B regions 1 and 2 shown in Figure 4E and F.

**Figure S5.**
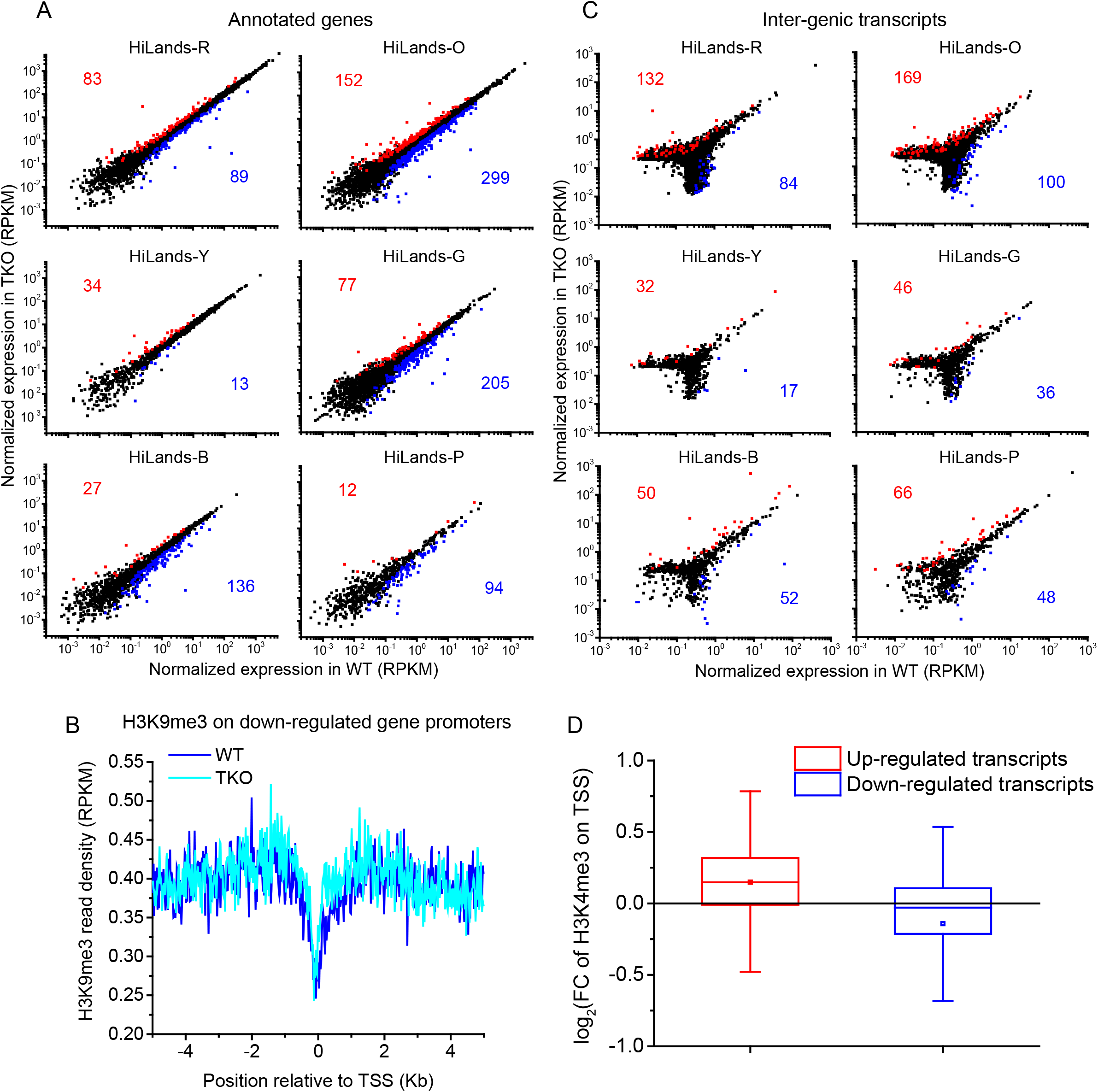
Transcription analyses (Support Figure 5) **A.** Scatter plots of annotated gene expression changes upon lamin loss in mESCs in 6 HiLands. Red dots and numbers represent up-regulated genes and blue dots and numbers represent down-regulated genes upon lamin loss called by edgeR software. Black dots are unchanged genes. **B**. Distribution of H3K9me3 reads within 5 Kb up- and down-stream of down-regulated TSS. **C.** Scatter plots for changed unannotated inter-genic transcripts upon lamin loss in mESCs in 6 HiLands. Red dots and numbers represent up-regulated genes and blue dots and numbers represent down-regulated genes upon lamin loss called by edgeR software. Black dots are unchanged genes. **D.** H3K4me3 changes at transcription start sites of up- and down-regulated transcripts from both the annotated genes and unannotated intergenic regions.

**Figure S6.**
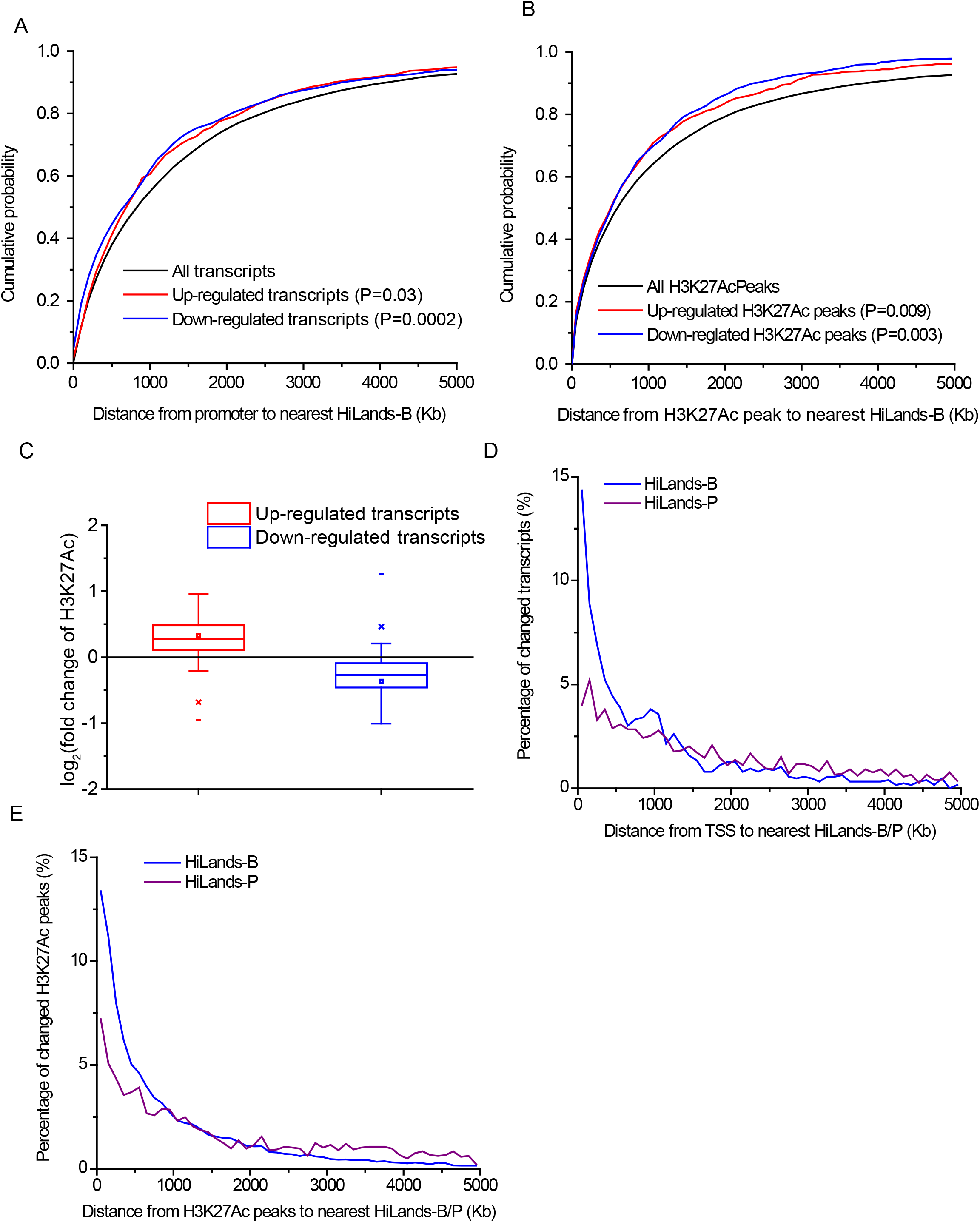
Analyses of transcription changes (support Figure 6) **A.** Cumulative probability curves for the distance distribution of up-regulated, down-regulated, and all TSS (annotated genes and intergenic transcripts) to the nearest HiLands-B. Only genes in HiLands-R, -O, -Y, and -G are used. P-value are from Kolmogorov-Smirnov test. **B.** Cumulative probability curves for the distance distribution of up-regulated, down-regulated, and all H3K27Ac peaks to the nearest HiLands-B. Only peaks in HiLands-R, -O, -Y, and -G are used. P-value are from Kolmogorov-Smirnov test. **C.** Change of H3K27Ac near the TSS of up- and down-regulated transcripts from both the annotated genes and unannotated intergenic regions. **D-E.** A plot showing the distance distribution of transcription start sites (TSS) of altered transcripts (D) or altered H3K27Ac peaks (E) upon lamin loss to the borders of the nearest HiLands-B and P. Only the TSS and H3K27Ac in HiLands-R, -O, -Y and -G are plotted.

## Supplementary Table Legends

**Table S1.** Statistics of our WT and TKO Hi-C libraries.

**Table S2**. Lists of TAD boundaries in WT (sheet 1) and lamin-TKO (sheet 2) and the change of TAD-TAD interactions (sheet 3).

**Table S3**. Lists of primers used to amplify the Oligopaint libraries for FISH probe production (sheet 1) and for 4C studies (sheet 2).

**Table S4**. Mean emerin-DamID values in 100-Kb windows along the genome in WT and lamin-B DKO mESCs treated by lamin-A/C RNAi (equivalent to lamin TKO) mESCs. The proportion of each 100-Kb window overlapping with different HiLands are also shown.

**Table S5.** Comparisons of H3K27me3 (sheet 1), H3K9me3 (sheet 2), H3K4me3 (sheet 3), H3K4me1 (sheet 4), and H3K27Ac (sheet 5) between our WT and lamin TKO mESCs.

**Table S6.** RNA-seq differential expression analysis for annotated genes (sheet 1) and unannotated intergenic transcripts (sheet 2) between our WT and lamin TKO mESCs. The location of each promoter with respect to HiLands are shown. The GO-terms of the down-regulated genes upon lamin loss in mESCs are shown in the third sheet. Sheet 4 shows the genes and their expression change in HiLands-B that showed detachment from the nuclear periphery consistently from both emerin Dam-ID and C-score data.

## STAR*METHODS

**Key Resources Table**

**Antibodies**

**Chemicals**

**Reagent**

**Critical commercial kits**

**Experimental models: cell lines**

**See Table S3 for primer sequences used for making oligo probes for FISH and for 4C.**

**Software and algorithms**

**Contact for reagent and resource sharing**

Please contact the corresponding author Yixian Zheng for further information, reagents, and tools.

## METHOD DETAILS

### Cell lines and Culture Conditions

The derivation of wild-type and lamin-TKO mESCs were described in our previous publication (Kim et al., 2013b). WT and TKO mESCs were cultured using the same method as described (Kim et al., 2013b) for Hi-C, ChIP-seq, and RNA-seq experiments. For FISH, mESC stocks, which were prepared from the mESCs expanded on feeder cells, were thawed and grown using 0.1% gelatin-coated plates in the absence of feeder cells with KO-DMEM medium (Invitrogen) containing 15% fetal bovine serum (FBS, Invitrogen), 100 μM β-mercaptoethanol (Invitrogen), 2 mM Glutamax, 0.1 mM nonessential amino acids, 50 μg/ml penicillin/streptomycin, and 1,000 units/ml LIF (Millipore). When the cells became ~70% confluent, they were passaged to a new plate and incubated for 30 min at 37°C in 5% CO2 to allow the attachment of the residual feeder cells to the bottom of the plate. The supernatant containing mESCs were used for FISH.

### 3D FISH, mESC FACS, and karyotyping

#### Oligopaint probe design

We used the OligoArray2.1 software to select oligo probes for genomic regions. First, the mouse genome sequence of the selected chromosomes was segmented into 1-kb regions and combined into a FASTA file. Then we ran the OligoArray2.1 software using the parameters: -n 3 -N 3 -l 50 -L 50 -D 1000 -t 85 -T 99 -s 70 -x 70 -p 35 -P 80 -m "GGGG;CCCC;TTTTT;AAAAA" on the Memex super computer cluster at the Carnegie Institution for Science. Next, we adjusted the probe numbers to obtain about 3-fold higher probe densities for selected HiLands-B and -P regions than the probe density of the rest of the chromosome. We then randomly selected as probes the genomic sequences that matched the desired probe numbers for each sub-library region and the total synthesis limit of 92918 probes for each chromosome library pool. Finally, we selected primer pairs for each sub-library and common primer pairs for oligo labeling (Table S2). These primers, which do not pair with the genomic probe sequences used or with one another, were attached to either end of the genomic probes to construct the Oligopaint library.

#### Oligopaint library synthesis and probe preparation

The Oligopaint libraries were synthesized by CustomArray. Each library was in 80 μl TE buffer with concentrations of 157.9 ng/μl, 101.4ng/μl, 112.5 ng/μl and 89.3 ng/μl for chr1, chr4, chr13 and chr14, respectively. We largely followed the published method (Beliveau et al., 2012) for FISH probe preparation with some modifications. For HiLands-B and -P sub-library probe preparation, each sub-library was first amplified using 0.5 μl of the corresponding chromosome library pool and the sub-library specific PCR primers (synthesized by Eurofins). The PCR product was purified using a Qiagen PCR purification kit. 30-μl elution buffer was used to elute the amplified DNA with a final DNA concentration between 10-30 ng/μl. 0.5 μl of this PCR product was further amplified using the common primer pair with the 5’ of the forward primer labeled by either Alexa-488 or Texas Red and the 5’ of the reverse primer phosphorylated (both synthesized by IDT).

The second PCR products were purified using Zymo DNA clean & Concentrator-500 columns (Zymo Research, D4032) and then treated with lambda exonuclease (NEB, M0262L) to remove the unlabeled strand and to produce the single strand DNA probes (ssDNA). The ssDNA was precipitated with NH_4_OAc and 100% ethanol, washed with 70% ethanol, and dissolved in double distilled water. For the whole chromosome probe preparation, only one round of PCR was performed using the individual chromosome library pool as the template and the common primer pair that were labeled with fluorophore and phosphorylated, followed by purification, lambda exonuclease digestion, and precipitation as described above. The probe concentration was measured using a Nanodrop 2000 (Thermo Scientific). The yield was 2000-3000 pmol for a 10 ml reaction. The probes sizes (88bp) were verified by gel electrophoresis. We used 40 pmol per slide for FISH.

#### Oligopaint FISH of mESCs

Superfrost Plus VWR Micro Slides (VWR, 48311-703) were rinsed in 100% ethanol, air dried, and then coated with 0.1% gelatin (Millipore, ES-006-B) at 37°C for 30 min. 100 μl of a solution containing 1-2x10^6^ mESCs/ml was spotted on each slide coated with gelatin. The mESCs were allowed to adhere for ~3 h at 37°C. Slides were then rinsed with 1x PBS at room temperature (RT), then fixed at RT for 10 min in 4% paraformaldehyde (PFA) freshly diluted with 1x PBS from the stock 16% PFA (Electron Microscopy Sciences, 15710). For postfixation, slides were rinsed in 1x PBS, permeabilized in 0.5% Triton X-100 for 10 min at RT and rinsed briefly in 1x PBS. Then the slides were incubated in 20% glycerol (diluted with 1x PBS) for a minimum of 60 min at RT. Slides were frozen in liquid nitrogen for ~15 s and thawed at RT. This freeze and thaw was repeated 4x. The slides were then washed in 1x PBS for 3 times, 5 min each time, followed by a 5 min incubation in 0.1M HCl and 3 washes (1 min each) in 2x SSC (0.3 M NaCl, 0.03 M NaCitrate). Slides were then warmed to 37°C in 2xSSC + 50% formamide for ~4 h. After warming up, the hybridization cocktail containing the probes prepared as above were added onto the slides, followed by denaturation at 78°C for 3 min on a water-immersed heat block and hybridized at 37°C for at least 14 h in a humidified chamber. The slides were then washed: (1) for whole chromosome FISH: wash twice in 2x SSCT (2x SSC + 0.1% Tween-20) at 60°C for 15 min or (2) for HiLands-B or -P FISH: wash twice in 50% formamide for 30 min at 37°C with shaking, wash once in 20% formamide for 10 min at RT with rotation, followed by 4 quick washes with 2x SSCT. After these washes, the slides were further washed with 0.2x SSC at RT for 10 min, followed by DNA staining with DAPI (diluted in 0.2x SSC) at RT for 5 min. Excess DAPI was washed away. The area containing the mESCs was mounted with ProLong Diamond Antifade Mountant (Life Technologies).

#### Combining immunostaining of emerin with FISH

After the wash steps with formamide, the slides were dried and the area containing the mESCs was marked using a PAP-pen (Abcam, ab2601). The mESCs were blocked with 4% BSA in 1x SSCT for 30 min at RT, followed by incubation at RT for 2 h with emerin antibody (Santa Cruz, sc-15378) diluted (1:300) in1x SSCT containing 1% BSA. After incubation, the slides were washed 3 times for 5 min each in 1x SSCT and incubated with secondary antibody (Alexa 594 goat anti-rabbit, Invitrogen) diluted (1:1000) in 1x SSCT containing 1% BSA, at RT for 1 h. The excess antibody was washed away 3 times for 5 min each in 1x SSCT. The DNA was stained and the slides were mounted as above.

#### Confocal image acquisition for 3D FISH

Confocal images were acquired using a Leica SP5 confocal microscope equipped with a 63x/1.4 objective and an electron multiplying charge-coupled device camera. For each set of experiments, images were acquired as confocal stacks at 126 nm per step in the z-axis using the same laser setting. 3D images were constructed from the stacks after correcting the channel shift using the FocalCheck fluorescence microscope test slide #1 (Molecular Probes).

#### Quantification of 3D FISH

The Imaris software (Bitplane) was used for all the quantifications. For double blind analyses, one person coded each set of 3D FISH image with a number corresponding to its true feature. These images were then randomized and coded with another set of numbers. These randomized images were given to another person to perform the quantification. For each FISH experiment, at least 50 nuclei were quantified. To measure the volume and surface area of selected HiLands-B, -P, or whole chromosomes, the surfaces were created based on the 3D FISH images. The surfaces were not smoothed. The minimal threshold was determined manually by adjusting the value until the surfaces just covered the entire FISH signal, while the maximum threshold was automatically determined by the software. The surfaces with greater than 10 voxels were then created. If the FISH signal was discontinuous, several surfaces would be generated for one HiLands region or chromosome. In this case the volume and surface area will be the sum of the discontinuous regions.

To measure the distance from the HiLands-B regions (Oligopaint signal, Alexa-488) to the nuclear periphery labeled by the emerin antibody (Alexa-594), each FISH signal was identified using the ‘Spots object’ function in Imaris. The diameter of the spot that identified the FISH signal was manually adjusted so that each spot recognized one HiLands-B FISH signal and covered most of its volume. The ‘Cells object’ in Imaris was used to identify the nuclear envelope labeled by the emerin antibody. The Cell type was chosen as Cell membrane; The smallest diameter of the Cell was adjusted manually to ensure that the emerin signal was represented as a solid shell enclosing or right on the FISH spots. Due to the harsh FISH conditions, a fraction of the nuclei exhibited extensive distortion such that the emerin staining appeared highly irregular with extensive invaginations. In these cases, the emerin signal could not form a solid shell that either overlapped or enclosed the FISH spots by adjusting the Cell smallest diameter. These nuclei were not included in the quantification. The FISH spots were imported into the Cell as vesicles. The distance from the center of the FISH spot to the nearest Cell shell was measured as the distance between the HiLands-B region and the nuclear envelope. The measured distance was automatically exported as "vesicles import distance to cell membrane" by Imaris. If the FISH spot overlapped with the Cell shell surface but its center was outside of the surface, it cannot be imported into the cells and the distance for the spot was regarded as 0.

#### FACS analyses of the cell cycle of lamin TKO and WT mESCs

FACS analyses of the cell cycle were performed as described previously (Pozarowski and Darzynkiewicz, 2004) with modification. Briefly, the cells were harvested, washed with 2% FBS diluted in 1x PBS and then fixed in 70% ice-cold ethanol at 4°C for at least 1h. The fixed cells were washed twice with ice-cold 1x PBS, followed by incubation in propidium iodide (PI, Sigma, P 4170) staining solution (25 μg/ml PI and 0.5 μg/ml RNase A in 1x PBS) at 4°C for at least 4 h. After staining, the cells were analyzed by FACS.

#### Karyotyping of lamin TKO and WT mESCs

Actively growing mESCs were incubated in 50 ng/ml colcemid solution for 2 h to accumulate mitotic cells. After colcemid treatment, the cells were harvested with trypsin, resuspended in fresh medium and treated with 75 mM KCl solution at 37°C for 15min. Then the cells were fixed in cold methanol and glacial acetic acid (3:1) on ice for 10 min. After fixation, the cells were dropped onto a pre-wetted slide and air-dried. Finally, the slides were stained with 1 μg/ml Hoescht 33258, washed and mounted in Immu-Mount (Thermo Scientific). The chromosomes were counted using a 100X oil immersion objective on a Nikon Eclipse E800 microscope.

### Hi-C library preparation and sequencing

#### Cell harvest and crosslinking

WT or TKO mESCs were cultured in feeder-free conditions as described (Kim et al., 2011). Five to eight million feeder-free mESCs were harvested and resuspended in 10 ml basal culture medium [10% FBS, 100 μM β-mercaptoethanol (Life Technologies), 2 mM L-glutamine, 0.1 mM nonessential amino acids, 1 mM sodium pyruvate, and 50 μg/ml penicillin/streptomycin in GMEM (Life Technologies)]. Cells were fixed with 1% formaldehyde for 10 min at room temperature. After fixation, formaldehyde was quenched by adding 125 mM glycine. The cells were incubated at RT for 5 min, followed by an additional incubation on ice for 15 min. The cells were centrifuged at 1,000g for 10 min, and the cell pellets were washed with ice-cold 1x PBS.

#### Cell lysis and the first enzyme digestion

The crosslinked cell pellet was resuspended and lysed in Hi-C lysis buffer [10mMTris, pH8.0, 10 mM NaCl; 0.2% IGEPAL CA-630 (NP-40),1x protease inhibitor cocktail (Roche, 04693132001)], rotating at 4°C for 30 min. The lysed cells were centrifuged at 1,000 g for 5 min. The pellet was washed with 1ml ice-cold 1.2x NEBuffer 3.1 (NEB, B7203S), and then resuspended in 0.4 ml 1.2x NEBuffer 3.1. To remove uncrosslinked proteins, 6 μl 20% SDS were added, followed by incubation at 65°C for 10 min. 40 μl 20% Triton X-100 were added to quench SDS and mixed carefully by inverting several times. The suspension was incubated at 37°C for 10 min. To digest the crosslinked chromatin, 30 μl 50 units/μl BglII enzyme (NEB, R0144M) were added to the suspension. The enzyme reactant was incubated overnight at 37°C.

#### Biotin filling and proximity ligation

To mark the cohesive ends generated by BglII with biotin, 1.5 μl 10 mM dATP, 1.5 μl 10 mM dGTP, 1.5 μl 10 mM dTTP (Qiagen), 3.0 μl 5 mM biotin-16-dCTP (Axxora, JBS-NU-809-BIO16) and 10 μl 5 units/μl Klenow (NEB, M0210L) were added. The mixture was incubated at 37°C for 45 min, and then placed on ice. The blunted chromatin DNA was ligated by adding 750 μl 10x T4 ligase buffer (NEB, B0202S), 75 μl 10 mg/ml BSA (NEB), 6151.5 μl distilled water, and 30 μl 30 units/μl T4 DNA ligase (Thermo Scientific, EL0013). A total of 7.5 ml ligation mixture was incubated overnight at 16°C.

#### Decrosslinking and DNA purification

To reverse crosslinking, 25 μl 20 mg/ml proteinase K (Life Technologies, 25530-049) was added and incubated overnight at 65°C. After cooling at RT, 10 ml phenol (pH 8.0) were added to the suspension, and vigorously mixed by vortexing for 2 min. The suspension was centrifuged for 5 min at 3,500 rpm with a standard tabletop centrifuge. The aqueous phase was mixed with 10 ml phenol:chloroform (1:1). After centrifugation, the supernatant was transferred to a 50 ml Oak Ridge centrifuge tube (Nalgene) and the total volume was brought to 10 ml with 1x TE (pH 8.0). To precipitate the ligated DNA, 1 ml 3M sodium acetate, 5 μl 15 mg/ml GlycoBlue (Life Tech, AM9515), and 10 ml isopropanol were added, and mixed by inverting. The mixture was incubated at -80°C for at least 1 h. The suspension was centrifuged at 38,000 g, 4°C for 30 min. The precipitated pellet was resuspended in 450 μl 1x TE (pH 8) and the suspension was transferred to an 1.5 ml Eppendorf tube. DNA was extracted with 500 μl phenol:chloroform (1:1). After phenol:chloroform extraction, 40 μl 3M sodium acetate, 1 μl GlycoBlue, and 1 ml 100% ethanol were added to the supernatant, and incubated at -80°C for 30 min. To precipitate DNA, the suspension was centrifuged at 18,000 g for 20 min. The pellet was washed with 70% ethanol and then air dried completely. The dried pellet was resuspended in 45 μl 10 mM Tris (pH 8.0). DNA concentration was determined using a Quant-iTdsDNA assay kit (Life Tech, 33120). 5 μg DNA was used for the next step.

#### Removing biotin from un-ligated DNA

5 μg DNA was mixed with 10 μl 10x NEBuffer 2.1, 12.5 μl 10 mM dNTP mix (Qiagen, 201900), 1.6 μl 3 units/μl T4 DNA polymerase (NEB, M0203S), and the total reaction volume was brought to 100 μl with distilled water. The mixture was incubated at 12ºC for 2 h. The 3’-5’ exonuclease activity of the T4 DNA polymerase degraded the un-ligated biotinylated DNA. The reaction was stopped by adding 2 μl 0.5 M EDTA. DNA was purified with a QIAquick PCR purification kit (Qiagen), eluting with 50 μl 10 mM Tris (pH 8.0).

#### Streptavidin capture and the second enzyme digestion

To bind biotinylated DNA to streptavidin beads, 30 μl Dynabeads MyOne Streptavidin C1 (Life Technologies, 65501) were washed and resuspended in 50 μl 2x binding buffer (10 mM Tris, pH 8.0, 0.1 mM EDTA, 2 M NaCl), 50 μl biotinylated DNA was added, and the mixture was incubated at RT for 30 min, constantly mixing with an Intelli mixer (Cole Parmer, UX-51202-22) or an equivalent. After washing with 1x binding buffer (10 mM Tris, pH 8.0, 0.1 mM EDTA, 1 M NaCl) and 1x NEBuffer 4.1, the beads were resuspended in 50 μl 1x NEBuffer 4.1. To trim biotinylated DNA into small fragments, 1μl 10 units/μl AluI (NEB, R01317S) was added and incubated at 37°C for 60 min. After AluI digestion, the beads were subsequently washed with 1x binding buffer and 10 mM Tris (pH 8.0), and resuspended in 16 μl 10 mM Tris (pH 8.0).

#### Paired-end sequencing library preparation

To ligate paired-end Illumina sequencing adaptors to the AluI digested DNA fragments, beads were processed with NEBNext^®^ DNA Library Prep Reagent Set (NEB, E6000S) as follows: For dA-tailing, the beads in 16 μl 10 mM Tris (pH 8.0) were mixed with 2.5 μl 10x NEBuffer 2, 5.0 μl dATP, 1.5 μl Klenow (all included in the NEB kit), and incubated at 37°C for 30 min; constant agitation is not required hereafter. dA-tailed beads were washed with 1x binding buffer, and resuspended in 5.0 μl 10 mM Tris (pH 8.0). To ligate sequencing adaptors, the beads in 5.0 μl 10 mM Tris (pH 8.0) were mixed with 12.5 μl ligation reaction buffer, 5.0 μl NEBNext adaptor, and 2.5 μl Quick T4 ligase (all included in the NEB kit), and incubated at 20°C for 15 min. To cleave the hairpin loop of NEBNext adaptors, 1.5 μl USER enzyme (included in the NEB kit) was added and incubated at 37°C for 15 min. The beads were washed with 1x binding buffer and resuspended in 10 μl of 10 mM Tris (pH 8.0). To amplify adaptor ligated DNA, the beads in 10 μl of 10 mM Tris (pH 8.0) were mixed with 1.25 μl Universal PCR primer, 12.5 μl PCR master mix (included in the NEB kit), 1.25 μl of NEBNext Indexed primer (NEB, E7500S), and the PCR was performed with the following program (98°C 30 sec, 8 cycles of [98°C 10 sec, 65°C 30 sec, 72°C 30 sec], and 72°C 5min). PCR amplified Hi-C library DNA was purified with AMPure XP magnetic beads (Beckman Coulter). The quality of Hi-C library DNA was assessed with a Bioanalyzer (Agilent, 2100). Successful Hi-C library DNA ranged from 200bp to 1kb in size with a clear single peak at about 600 bp. Paired-end sequencing was performed with a HiSeq 2000 (Illumina) for replicate 1 (WT and TKO, 50-bp paired-end) and Nextseq500 (Illumina) for replicate 2 (WT and TKO, 75-bp paired-end).

### RNA-sequencing

Total RNA was extracted from WT and TKO ESC using RNeasy Plus Mini Kit (Qiagen, 74134). Ribosomal RNA was removed with Ribo-Zero rRNA Removal Kit (Illumina, MRZH11124). Sequencing libraries were built using TruSeq Stranded mRNA Library Prep Kit (Illumina, RS-122-2102).

### ChIP-sequencing

4 × 10^6^ TKO or WT mESCs were harvested and fixed as described in the Hi-C method section. The crosslinked cells were resuspended in *Nucleus*/Chromatin Preparation (NCP) buffer I [10 mM Hepes, pH 6.5, 10 mM EDTA, 0.5 mM EGTA, 0.25% Triton X-100 and 1x protease inhibitors (Roche)] and incubated at 4°C for 10 min with gentle rotation. The suspension was centrifuged at 2,000g for 5 min. The pellets were resuspended in NCP buffer II (10 mM Hepes, pH 6.5, 200 mM NaCl, 1 mM EDTA, 0.5 mM EGTA and 1x protease inhibitors), followed by centrifugation. The pellets were resuspended in SDS lysis buffer (10 mM Tris, pH 8.0, 1 mM EDTA, 1% SDS and 1x protease inhibitors). The suspension was sonicated with a Bioruptor sonicator (Diagenode) to break chromatin DNA into about 200 bp in size. The sonicated DNA was diluted 10fold with ChIP dilution buffer (0.01% SDS, 1.1% Triton X-100, 1.2 mM EDTA, 16.7 mM Tris, pH 8.1, and 167 mM NaCl). To precipitate chromatin, 40 μl protein A-Dynabeads (Life Technologies, 10001D) resuspended in 0.5 ml ChIP binding buffer (0.01% Tween-20 in 1x PBS) were incubated with 4 μg of antibodies for each IP at 4°C for 6 h. After washing with ChIP binding buffer, the beads were incubated with diluted sonicated chromatin DNA suspension overnight, continuously rotating. The beads were sequentially washed with low salt wash buffer (20 mM Tris pH 8.1, 0.1% SDS, 1% Triton X-100, 2 mM EDTA, and 150 mM NaCl), high salt wash buffer (20 mM Tris pH 8.1, 0.1% SDS, 1% Triton X-100, 2 mM EDTA, and 500 mM NaCl), LiCl buffer (10 mM Tris, pH8.0, 250 mM LiCl, 1% IGEPAL CA630 (NP40), 1% Sodium deoxycholate, and 1 mM EDTA), and TE buffer. To reverse crosslinking, the beads were incubated with elution buffer (10 mM Tris, pH 8.0, 1% SDS, 1 mM EDTA and 0.2 µg/μl Proteinase K) at 65°C overnight. Supernatant was collected, extracted with Phenol:Chloroform (1:1), and precipitated with ethanol. The pellets were washed with 75% ethanol, air-dried, and resuspended in 10 mM Tris (pH 8.0). DNA concentration was determined with a Quant-iTDS DNA kit (Life Tech, Q-33120). 1 ~ 400 ng of DNA were used to prepare ChIP-seq libraries with a TruSeq DNA prep kit (Illumina) according to the manufacturer’s recommendation. Single-end 50-bp sequencing was performed with a HiSeq 2000 (Illumina). Antibodies used for ChIP were anti-H3K4me1 (Abcam, ab8895), anti-H3K4me3 (Cell Signaling, 9751), anti-H3K9me3 (Abcam, ab8898), anti-H3K27ac (Abcam, ab4729), and anti-H3K27me3 (Millipore, 07-449).

### 4C-sequencing

4C libraries were prepared as previously described with minor modification (van de Werken et al., 2012). Briefly, 1×10^7^ cells were crosslinked with 2% formaldehyde at RT for 10 min, followed by glycine quenching, cell lysis, DpnII digestion, and T4 ligation. The ligation product was reverse-crosslinked and then further trimmed with Csp6I. The resulting DNA was circularized by T4 ligation. The second ligation product was purified with DNA Clean & Concentrator™-100 (Zymo Research, 4029). 4C libraries were amplified from the purified second ligation product with 4C primers as specified in Supplementary Table S3. The primers were designed using the following website: http://mnlab.uchicago.edu/4Cpd/.

### Hi-C data analyses

#### Hi-C data preprocessing

Hi-C reads were mapped and filtered using HiC-Pro (Servant et al., 2015). We then performed additional filtering to remove reads in the Encode blacklist (Consortium, 2012). A raw contact matrix was generated for each sample at 20-Kb resolution by counting the number of reads between each pairs of 20-Kb bins, and combined raw contact matrices were generated for WT and TKO by adding the matrices of replicates.

#### Hi-C data normalization

We performed Hi-C data normalization using the Iterative Correction and Eigenvector decomposition (ICE) method. We first filtered Hi-C windows with low or extremely high coverage (<1/5 or >5 fold of the median coverage) in the raw contact matrices generated above, then we ran the ICED script in the HiC-Pro package (Servant et al., 2015) to get the normalized contact matrices.

#### TAD inference

We used the Insulation Score (IS) method (Crane et al., 2015) to identify TADs boundaries. For each 20-Kb window, the total ICE-normalized interaction across the window from 500-Kb up- and down-stream of the window were added up. Then this number of total interaction was further normalized by its genomic average and log_2_ transformed to get the IS. Local minima of the IS indicate TAD boundaries (Crane et al., 2015).

We used the same process and criteria as (Crane et al., 2015) to identify the TAD boundaries, except that we added an additional criterion: IS<0, to ensure that the interaction across TAD boundaries are lower than the genomic average.

#### Hi-C data visualization

For visualization of the Hi-C data, as shown in Fig. 1A-B and 2D, we used the dynamic binning method as used in the HiFive package (Sauria et al., 2015) to account for sparse coverage. A bin pair was expanded in all directions (not crossing the diagonal) until the number of reads in the bin pair reached a minimum number (20 in our case). Then the contact frequency of the bin pair was calculated by the average contact frequency in the expanded bin pair. In this way, the bin pairs of high interactions retain the high resolution, while the bin pairs of low interactions avoid high noise.

For visualization of the Hi-C interaction changes, as shown in Fig. 1E and 6F-G, we used a similar dynamic binning method. We required the expanded bin pair to have no less than the minimum number of 20 reads in both WT and TKO datasets. Then the log_2_ fold change was calculated based on the expanded bin pair.

#### C-score model and estimation

As described in the main text, we allowed each individual cell to have its own A-B compartment segregation, and each specific genomic window (denoted by *i*) has a chance *P*_*i*_ to be in the B-compartment in an individual cell.

Considering two genomic window *i* and *j*, if their distribution in A and B compartments are independent of each other, the chance that they are in the same compartment in an individual cell should be

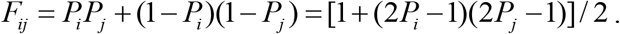

We define the C-score *C*_*i*_ = 2*P*_*i*_ −1, which is ranged between -1 and 1, and get *F*_*ij*_ =(1 + *C*_*i*_*C*_*j*_)/2.

Since in individual cell, windows in different compartments are unlikely interacting with each other, the number of Hi-C contact between two genomic window *i* and *j* should be proportional to *F*_*ij*_. Further considering the effect of linear genomic distance and experimental biases, we get *E*_*ij*_ = *B*_*i*_ *B*_*j*_ *H*(*d*_*ij*_)(1 + *C*_*i*_*C*_*j*_), where *E*_*ij*_ is the number of expected contact between windows *i* and *j*, *d*_*ij*_ is the linear distance, *H*(*d*_*ij*_) is the function of the distance dependency, *B*_*i*_ and *B*_*j*_ are the bias factors from Hi-C experiment. We assume the observed number of contact (*n*_*ij*_) follows a Poisson distribution *n*_*ij*_ ~ *Poisson*(*E*_*ij*_), and get the log-likelihood function from the Hi-C data.

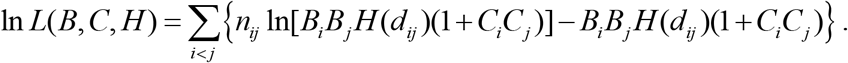

Here since we assumed the distribution of window *i* and *j* in A and B compartments are independent of each other, we used only window pairs with at least 1Mb distance. We also used only intra-chromosome interactions to reduce noise. We then used a maximum-likelihood approach to estimate all the parameters (B, C, and H).

Note that –ln *L* is a convex function for each individual parameter. Thus our maximum-likelihood estimation was done by iteratively optimizing each of the model parameter one by one, conditioning on other parameters, which ensures monotone increase of the likelihood function in each step. This monotone increase of the likelihood function ensures convergence based on the monotone convergence theorem. Details of the optimization steps are described as below:

For *B*_*i*_, we have 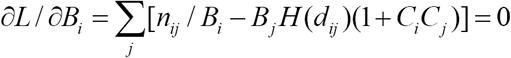, and 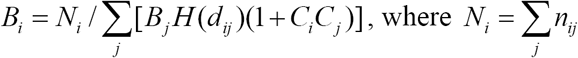.

For *H*, we have 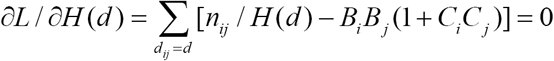 and 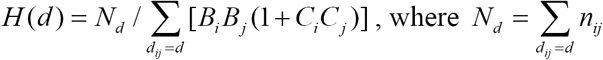.

Since *N*_*d*_ may be low for large *d,* we used exponentially growing distance bins [*d*_*n*_,*d*_*n+1*_), where *d*_0_ =1 Mb, each *d*_*n*+1_ = *d*_*n*_ × 10^0.04^, and all distances within each bin were regarded as equal.

For *C*_*i*_, we remove irrelevant parts in the log-likelihood function and get ln 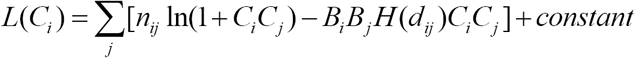. The maximum point of this function in (-1,1) does not have an explicit solution, but can be solved by Newton method, since –ln*L*is a convex function of *C*_*i*_.

The whole algorithm works as following: We first randomly generated the initial parameters, and then we performed iterative optimization for each parameter one by one as described above, in the order of *H*, *B* and *C*. The algorithm is considered convergent when the increase of the value of the log-likelihood function between two round (one round means updating all parameters once) is less than 1, and we can obtain the convergent values of C-scores. Each chromosome was analyzed separately, since interchromosome reads were not used. In practice, the algorithm converged in ~ 20 rounds, and was not sensitive to the initial parameters, since we tried 10 random initial parameters and obtained the same results. Note that if we flip the signs of all C-scores on one chromosome, the likelihood function will remain the same. Therefore, the final signs of C-scores on each chromosome was compared to lamin-B1 DamID, and the signs were globally flipped if positive C-scores generally corresponded to lamin-B1-negative regions.

### DamID data analysis

Two replicates of normalized emerin-damID data for WT mESC and lamin-B DKO mESCs treated by lamin-A/C RNAi (lamin-TKO equivalent) were downloaded from the GEO database (GSE62685). The DamID values for each probe was mapped back to the genome locations of mm9 according to the platform GPL8840. Average emerin-DamID data for both WT and lamin-depleted mESCs were calculated from the arithmetic mean of two biological replicates. Then for 100-Kb sliding windows along the genome, we calculated the mean emerin-DamID values from all probes within the 100Kb window as the emerin-DamID values for the window.

### RNA-seq data analysis

#### RNA-seq analysis for annotated genes

All RNA-seq samples (4 WT replicates and 4 TKO replicates) were sequenced on an Illumina NextSeq 500 using single-ended 75-bp mode. The resulting reads were mapped to the mouse genome mm9 using Tophat 2.1.0 (Kim et al., 2013a), and we used the GENCODE GTF file (vM1) as known transcripts (-G option). We used htseq-count (Anders et al., 2015) to count reads on genes defined by the GENCODE GTF file. Only genes in the “lincrna”, “protein coding”, and “processed transcript” types were considered.

The read counts were then processed using edgeR (Robinson et al., 2010). The read counts were normalized using the TMM normalization method in the edgeR package. Differentially expressed genes were called by edgeR using the GLMfit method, and batch effects were considered. The WT biological replicate 1-2 and lamin TKO biological replicate 1-2 were considered as one batch; the WT biological replicate 3-4 and lamin TKO biological replicate 3-4 were considered as another batch. We used fold change>1.5 and FDR<0.05 as threshold for differential expression.

#### RNA-seq analysis for unannotated inter-genic transcripts

The RNA-seq reads (same as above) were remapped using HISAT2 (Kim et al., 2015). We combined the mapped BAM files from different replicates and got one combined BAM files for WT and TKO. Then we ran Stringtie (Pertea et al., 2015) for de novo transcript assembly from the two BAM files and got two GTF file. The two GTF files were combined using Cuffmerge (Trapnell et al., 2010) based on the reference GTF file (GENCODE vM1) and genome sequence (-s option, with mm9 genome, for repeat region masking) to get a new merged GTF file. We then filtered the merged GTF file and only genes of class code “u” (intergenic transcripts not overlapping with annotated genes) were kept for further analysis.

We used htseq-count (Anders et al., 2015) to count reads on genes defined by the merged GTF file of intergenic transcripts. The differential expression analysis using read counts on intergenic transcripts was similar to the analysis for annotated genes described above using edgeR, except that the normalization was based on the number of reads mapped to annotated genes. We used fold change>1.5 and FDR<0.05 as threshold for differential expression.

### 4C-seq data analysis

4C data were processed using 4cseqpipe (van de Werken et al., 2012) to get normalized signal density. Then for comparison between WT and TKO, we calculated the fold change of signal density for each 20-Kb bin. To account for low signal at regions far from the bait, we expanded those bins (>500k from the bait) according to the distance to the bait: *bin size=0.04*×*distance*. The fold changes were calculated using the total signal density within the expanded bins.

### ChIP-seq data analyses

Chip-seq libraries were sequenced on an IlluminaHi-seq 2000 with single-end 50-bp mode. The reads were mapped to the mm9 genome using Bowtie 1.1.2 (Langmead et al., 2009) with "-v 2 -m 1" option.

For H3K4me3 mapping, reads were directly counted on promoters and the average fold change from 3 biological replicates was calculated for lamin TKO against WT.

For H3K27Ac, ChIP-seq reads for 3 biological replicates were first combined, then the peaks were called by Homer (Heinz et al., 2010) for the combined library. The parameters used were "-size 2000 -i Input_combined -L 0 -C 3 -minDist 2000 -tbp 3 ". Here "Input_combined" is the input library. Then the number of reads falling into each peak was counted for each replicate library of WT and lamin TKO. Differential peaks were called using edgeR GLMfit method similar to RNA-seq analysis. For H3K27Ac and RNA-seq correlation analysis, each H3K27Ac peak was assigned to the closest promoter.

For H3K4me1 (3 biological replicates), H3K27me3 (4 biological replicates), and H3K9me3 (3 biological replicates), the data processing was similar to H3K27Ac except that the peak calling parameters were " -style histone -i Input_combined -size 1000 -minDist 1000" to accommodate broader peaks. Then similarly the number of reads falling into each peak were counted for each library and differential peaks were called using edgeR.

### Data availability

All Hi-C, 4C, ChIP-seq, and RNA-seq data of WT and TKO mESCs in this study have been deposited in the GEO database (GSE89520).

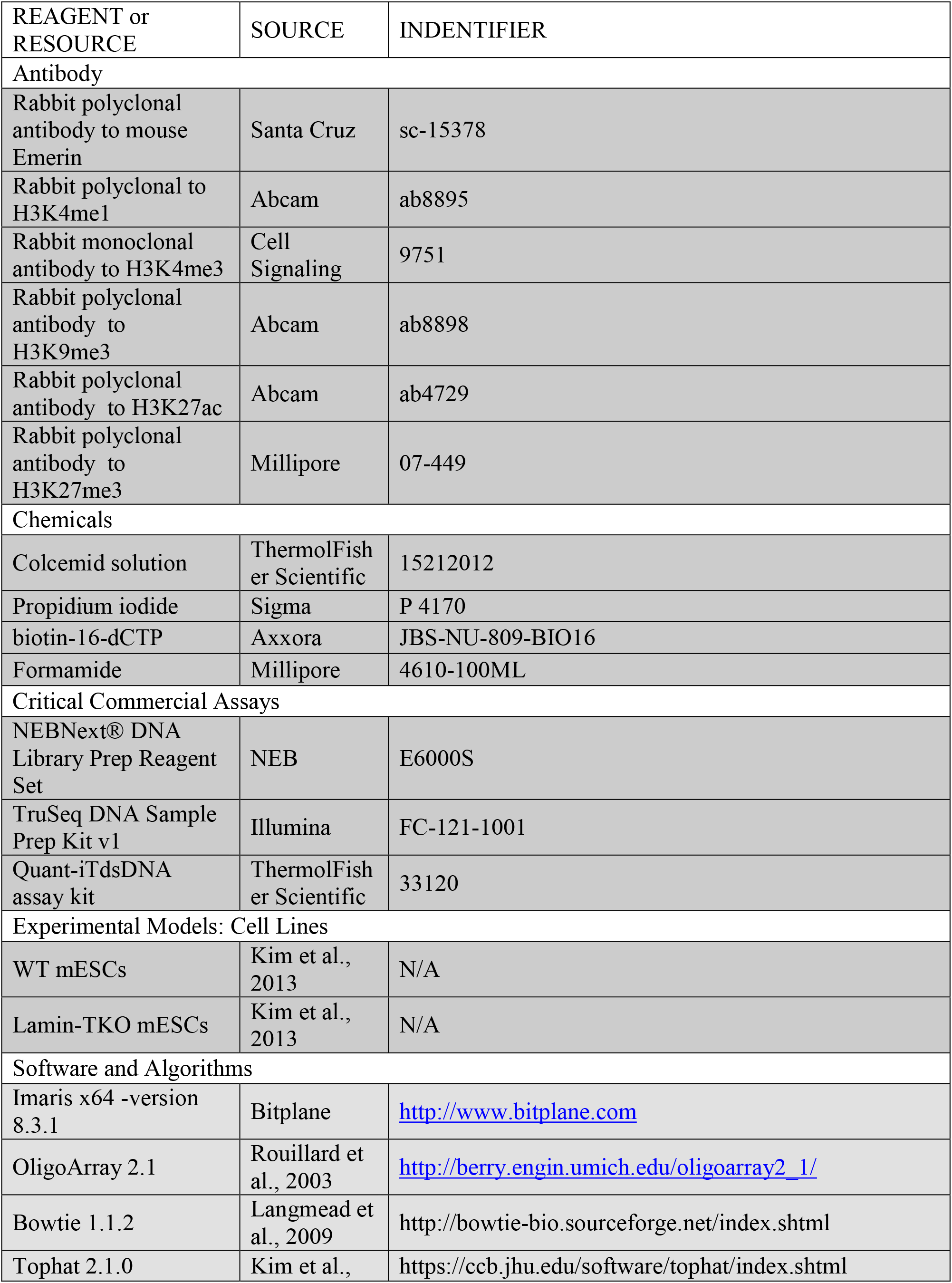
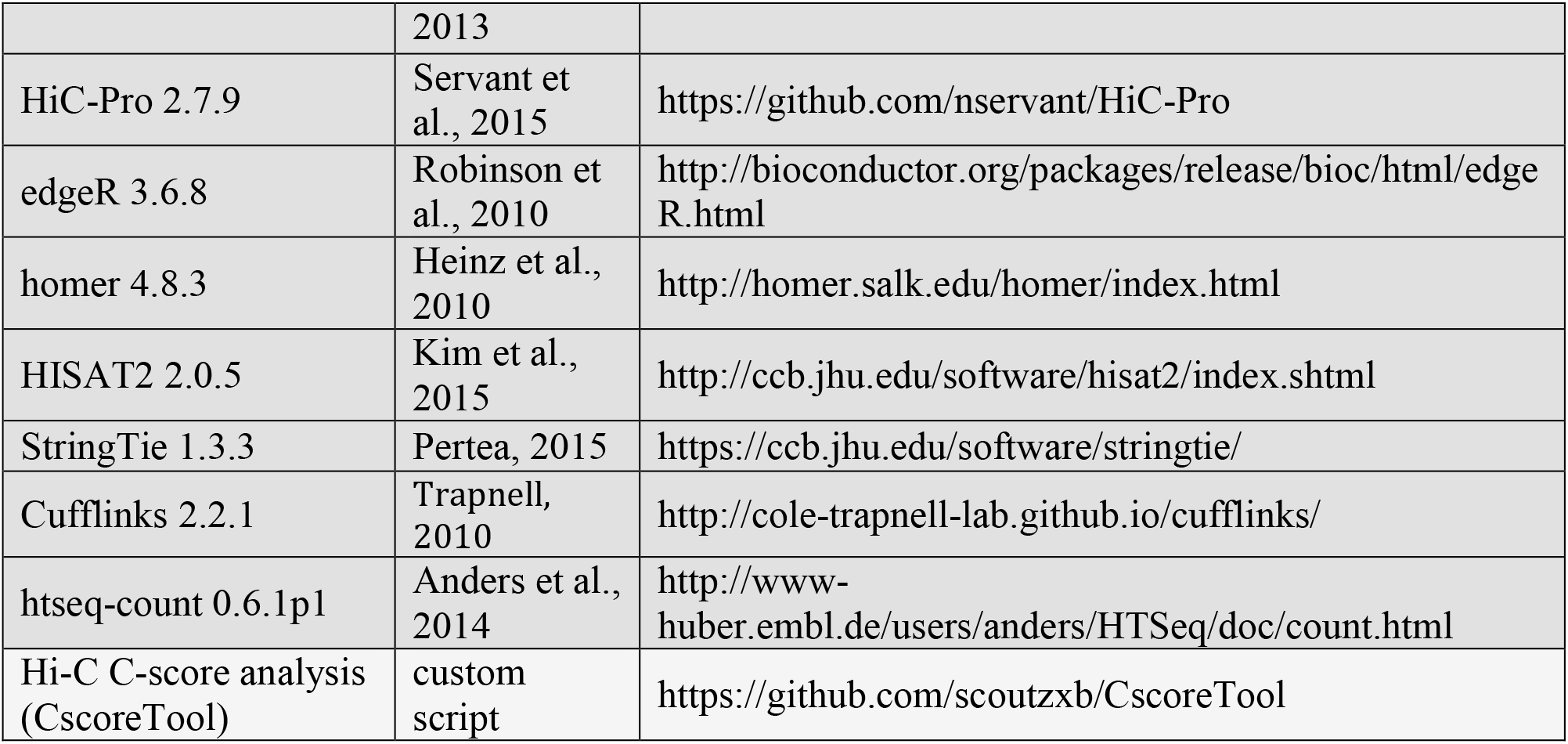

